# Heritable Large Deletions Using Type I-E CRISPR-Cas3 in Rice

**DOI:** 10.1101/2025.06.05.658196

**Authors:** Hiroaki Saika, Naho Hara, Shuhei Yasumoto, Toshiya Muranaka, Kazuto Yoshimi, Tomoji Mashimo, Seiichi Toki

## Abstract

Type I-E CRISPR-Cas3 derived from *Escherichia coli* (Eco CRISPR-Cas3) can introduce large deletions in target sites and is available for mammalian genome editing. The use of Eco CRISPR-Cas3 to plants is challenging because 7 CRISPR-Cas3 components (6 Cas proteins and CRISPR RNA) must be expressed simultaneously in plant cells. To date, application has been limited to maize protoplasts, and no mutant plants have been produced. In this study, we developed a genome editing system in rice using Eco CRISPR-Cas3 via *Agrobacterium*-mediated transformation. Deletions in the target gene were detected in 39–48% and 55–71% of calli transformed with 2 binary vectors carrying 7 expression cassettes of Eco CRISPR-Cas3 components and a compact all-in-one vector carrying 3 expression cassettes of Cas proteins fused to 2A self-cleavage peptide, respectively. The frequency of alleles lacking a region 7.0 kb upstream of the PAM sequence was estimated as 21–61% by quantifying copy number by droplet digital PCR, suggesting that mutant plants could be obtained with reasonably high frequency. Deletions were determined in plants regenerated from transformed calli and stably inherited to the progenies. Sequencing analysis showed that deletions of 0.1–7.2 kb were obtained, as reported previously in mammals. Interestingly, deletions separated by intervening fragments or with short insertion and inversion were also determined, suggesting the creation of novel alleles. Overall, Eco CRISPR-Cas3 could be a promising genome editing tool for gene knockout, gene deletion, and genome rearrangement in plants.

## Introduction

The sequence-specific nucleases of the CRISPR-Cas system are useful tools for genome editing in both basic research and molecular breeding in many organisms, including plants. Targeted mutagenesis using CRISPR-Cas9—the most widely used system—enables gene knockout and modification of gene expression. In addition, targeted mutagenesis using two or more guide RNAs (gRNAs) enables simultaneous knockout of multiple genes, introduction of larger deletions between gRNAs in a targeted promoter region, and generation of chromosome rearrangement including inversion and translocations (Gilbertson et al. 2025). CRISPR-Cas9 has also been applied to precise modification, such as base editors (Gaudelli et al. 2017; Komor et al. 2016; Nishida et al. 2016) and prime editing (Anzalone et al. 2019). CRISPR-Cas are divided into two classes: Class 2 CRISPR-Cas consists of a single Cas effector such as Cas9 and Cas12, whereas Class 1 CRISPR-Cas consists of multiple effector proteins. Class 1 Type I CRISPR-Cas induces DNA digestion in two steps: target sequence recognition by the complex of CRISPR-associated complex for antiviral defense (Cascade) with CRISPR RNA (crRNA), and subsequent DNA cleavage by Cas3 nuclease recruited by the Cascade complex. Class 1 CRISPR-Cas has been thought to be hard to use as a genome editing tool because of the requirement to express multiple CRISPR-Cas components in cells, although it is the major system in bacteria and archaea when compared with Class 2 CRISPR-Cas (Yoshimi and Mashimo 2022). Class 1 type I CRISPR-Cas is divided into 7 subtypes: I-A to I-G. Recently, the application of some Class 1 type I CRISPR-Cas systems to genome editing in eukaryotes has been reported. For example, genome editing in human cells was reported using type I-A (Hu et al. 2022), I-B and I-C (Tan et al. 2022), I-D (Osakabe et al. 2021), and I-E (Dolan et al. 2019; Morisaka et al. 2019). Many Class 1 CRISPR-Cas systems enable the introduction of large deletions (several kb), although their positions and sizes are not strictly determined. Class 1 CRISPR-Cas could be used to eliminate unnecessary genes, while both Cas9 and Cas12 enable the introduction of small insertions/deletions (indels) (Liu et al. 2022). In addition, Class 1 CRISPR-Cas has the potential to decrease off-target mutations due to longer spacers (32–37 nt) compared with Cas9 and Cas12 (Yoshimi and Mashimo 2022). Thus, Class 1 CRISPR-Cas has unique characteristics, and is expected to be an attractive genome editing tool also in plants. However, successful examples of genome editing using CRISPR-Cas3 in plants are limited to rice (type I-C, Li et al. (2023)), maize (type I-C and E, Li et al. (2023)), and tomato (type I-D named as TiD, Osakabe et al. (2020)). Osakabe et al. (2020) succeeded in genome editing in tomato using TiD—the type I-D CRISPR-Cas derived from *Microcystis aeruginosa*— and showed that these mutations were inherited to the next generation. The mutation patterns introduced by TiD were unique; both small indels (1–75 bp) and 2.5–7.3 kb of bidirectional and large deletions. In the TiD system, Cas10d has nuclease activity; Cas3d has no nuclease activity because it lacks a nuclease domain (Osakabe et al. 2020). The dual mutation patterns could be due to the characteristics of Cas10d nuclease (Wada et al. 2022). Li et al. (2023) applied Dvu I-C—a type I-C CRISPR-Cas derived from *Desulfovibrio vulgaris* str Hildenborough—to genome editing in rice and maize, and showed the genetic inheritance of deletions introduced by Dvu I-C. Type I-C Cascade complex is a relatively compact system and enables the introduction of unidirectional large deletions (Yoshimi and Mashimo 2022). To overcome the unpredictability of the size of deletions induced by Class 1 CRISPR-Cas, Li et al. (2023) established a system with the introduction of targeted large deletions using paired crRNAs; they also reported that the transient expression of Eco CRISPR-Cas3—a type I-E CRISPR-Cas derived from *E. coli*—induced large deletions by using paired crRNAs in maize protoplasts.

Type I-E CRISPR-Cas consists of 6 effector proteins: Cas5, Cas6, Cas7, Cas8, and Cas11 for the Cascade complex and Cas3 nuclease/helicase. With Eco CRISPR-Cas3, AAG, TAG, GAG, AGG, and ATG are recognized as a protospacer adjacent motif (PAM) sequence (Morisaka et al. 2019). After scanning the PAM sequence by EcoCas8 in the Cascade complex, an R-loop structure is formed by hybridizing the target DNA strand with crRNA. EcoCas3 is recruited at the R-loop structure and induces a nick in the non-target DNA strand. Double-stranded DNA is then unwound upstream of the PAM site by the helicase activity of EcoCas3. Target DNA is degraded by a combination of cleavage of the target strand in *trans* by collateral activity of EcoCas3 and repetitive cleavage of the non-target DNA strand in *cis*. As a result, long (0.5–80 kb), unidirectional deletions starting 0–400 bp upstream of the PAM sequence can be introduced (Yoshimi and Mashimo 2022). Successful applications of Eco CRISPR-Cas3 to genome editing have been reported in animal systems such as human cells, pig fibroblasts, and mice and rat zygotes (Miyagawa et al. 2022; Morisaka et al. 2019; Yoshimi et al. 2024). In mice and rat zygotes, DNA-free genome editing using mRNA and a ribonucleoprotein (RNP) complex of Eco CRISPR-Cas3 was reported (Yoshimi et al. 2024). In plants, however, to date there is only one report of Eco CRISPR-Cas3-mediated genome editing, which was in maize protoplasts, as described above (Li et al. 2023). As yet, there are no reports of the production of mutated plants by type I-E CRISPR-Cas. Besides targeted mutagenesis, type I-E CRISPR-Cas has been applied to transcriptional activators in maize immature embryos (Young et al. 2019) and random mutagenesis within long and defined regions in *Saccharomyces cerevisiae* (Zimmermann et al. 2023).

Here, we succeeded in producing regenerated plants with large deletions in a target gene using Eco CRISPR-Cas3 in rice by *Agrobacterium*-mediated transformation, and demonstrated that deletions confirmed in regenerated plants were inherited stably to the next generation. In addition, we determined the deletion frequency introduced by Eco CRISPR-Cas3 accurately using droplet digital PCR (ddPCR). The frequency of alleles lacking a region at 2.6 kb and 7.0 kb upstream of the PAM sequence in transformed calli was estimated at 21.1–60.8% and 39.5–74.1%, respectively, suggesting a reasonably high frequency of mutant regenerated plant production. These results showed the effectiveness of Eco CRISPR-Cas3-mediated genome editing in plants.

## Results and discussion

### Targeted mutagenesis by Eco CRISPR-Cas3 in rice calli

First, to investigate whether Eco CRISPR-Cas3 can be used for targeted mutagenesis in rice, we constructed binary vectors harboring expression cassettes of 6 proteins and crRNA (Fig. 1A). To achieve high and simultaneous expression in rice cells, the following strategy was employed: (1) codon optimization of Cas effectors with bipartite nuclear localization signals (bpNLS) to rice; (2) strong expression in rice calli using the maize ubiquitin promoter; (3) minimization of T-DNA length by separation of 6 Cas gene expression cassettes into 2 binary vectors: Cas537 (expressing EcoCas3, EcoCas5, and EcoCas7) and Cas6811 (expressing EcoCas6, EcoCas8, and EcoCas11); (4) higher DNA cleavage activity using crRNA consisting of pre-crRNA harboring an 86-nt leader sequence, and two 29-nt repeat sequences with an intervening 32-nt spacer sequence following a previous study in human cells (pre-crRNA (LRSR) in Morisaka et al. (2019)). To facilitate detection of genome editing events, *OsPDS* was selected as a target gene because knockout of *OsPDS* results in albino phenotype and makes it easier to distinguish genome-edited cells (Fig. 1B). In addition, a 32-nt spacer with 5′-AAG-3′ PAM was selected because AAG PAM showed the highest DNA cleavage activity in human cells (Morisaka et al. 2019). Rice calli were infected with a mixture of *Agrobacterium* harboring binary vectors Cas537 and Cas6811+OsPDS, and selected with hygromycin and G418 for 1.5 months. Some pieces of albino calli were found in transformed calli, suggesting that biallelic mutations had occurred in the *OsPDS* (Fig. 1C). To confirm whether deletions were introduced in *OsPDS* in transformed calli, PCR analysis was performed using primers designed to be longer upstream of the PAM because EcoCas3 introduces unidirectional deletion (Morisaka et al. 2019). Multiple fragments shorter than 8.7 kb were amplified in transformed calli, although there were no bands other than 8.7 kb corresponding to wild type in non-transformants (Fig. 1D). The frequency of transformed calli in which shorter amplicons were found was estimated as 39.4–48.2% (Supplementary Table S1). These results suggested that large deletion occurred specifically in *OsPDS* by Eco CRISPR-Cas3 with reasonably high frequency. Similar results were obtained in experiments using a different target designed in *OsDL* (Supplementary Fig. S1A-C).

**Figure 1.**
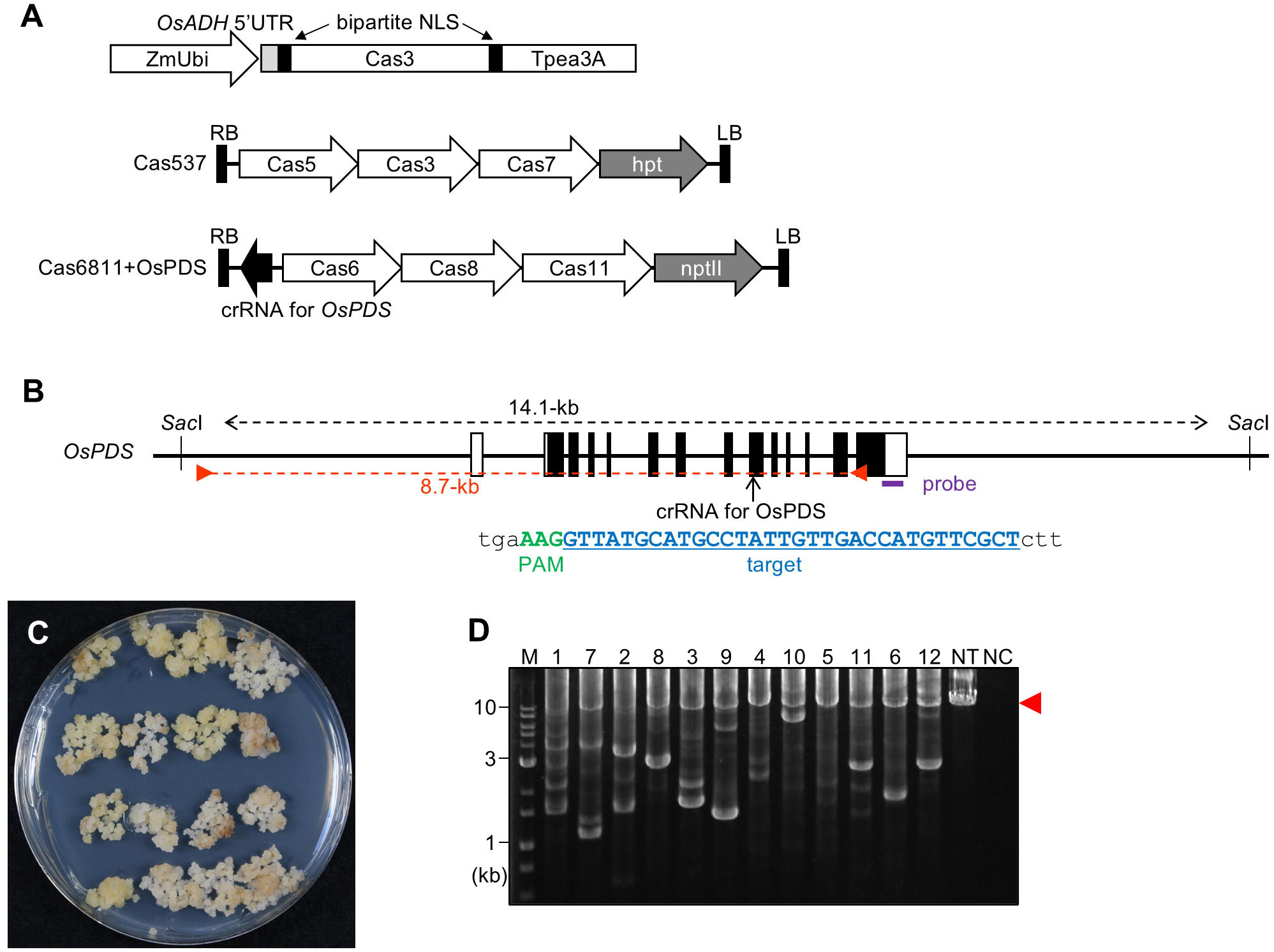
Eco CRISPR-Cas3-mediated targeted mutagenesis in rice calli. (A) Eco CRISPR-Cas3 vector targeting *OsPDS*. Top, EcoCas3 expression cassette. Grey and black shading show the 5′-untranslated region of rice alcohol dehydrogenase and nucleus localization signal (NLS), respectively. Middle, binary vector Cas537; bottom, binary vector Cas6811; ZmUbi; maize ubiquitin-1 promoter, Tpea3A; pea rbcs terminator, hpt; hygromycin phosphotransferase, nptII; neomycin phosphotransferase II, LB; left border. RB; right border. (B) Structure of *OsPDS* locus. Black and white boxes show coding regions and untranslated regions of *OsPDS*, respectively. Orange arrowheads and purple bar show PCR primers and the probe for Southern blot analysis, respectively. Green text, PAM; blue text, target sequence for Eco CRISPR-Cas3. (C) Rice calli transformed with binary vectors harboring Eco CRISPR-Cas3 with the crRNAs for *OsPDS* shown in A. (D) PCR analysis of transformed calli using primers shown in B. Red arrowhead, fragment size amplified in wild type. M; size marker, NC; negative control (no template), NT; non-transformant.

### Inheritance to progenies of mutations introduced by Eco CRISPR-Cas3

Hygromycin- and G418-resistant calli transformed with Cas537 and Cas6811+OsPDS were transferred to the regeneration medium. A total of 24 independent regenerated plantlets were obtained successfully from transformed calli and 12 plants showed albino phenotype (Supplementary Fig. S2A). Of these, 5 independent regenerated plants, including 3 lines of albino plants, were selected randomly and analyzed. No fragments and shorter fragments were found in albino plants #2 and #3, respectively, although there were no bands other than 8.7 kb corresponding to wild type in green plants #4 and #5 and non-transformants (Supplementary Fig. S2B). In addition, Sanger sequencing of PCR fragment #1 confirmed a 391-bp deletion showing that plants in which deletions were introduced in *OsPDS* had been obtained successfully. To confirm the genetic inheritance of mutations found in regenerated plants, we focused on *OsWx* encoding granule-bound starch synthase for amylose synthesis in endosperm, and designed targeted mutagenesis using Eco CRISPR-Cas3 (Fig. 2A). A total of 7 independent regenerated plants were obtained from rice calli transformed with Cas537 and Cas6811+OsWx. A large deletion was found in one line by PCR analysis (Fig. 2B). A total of 12 T1 seeds were obtained from this regenerated plant.

**Figure 2.**
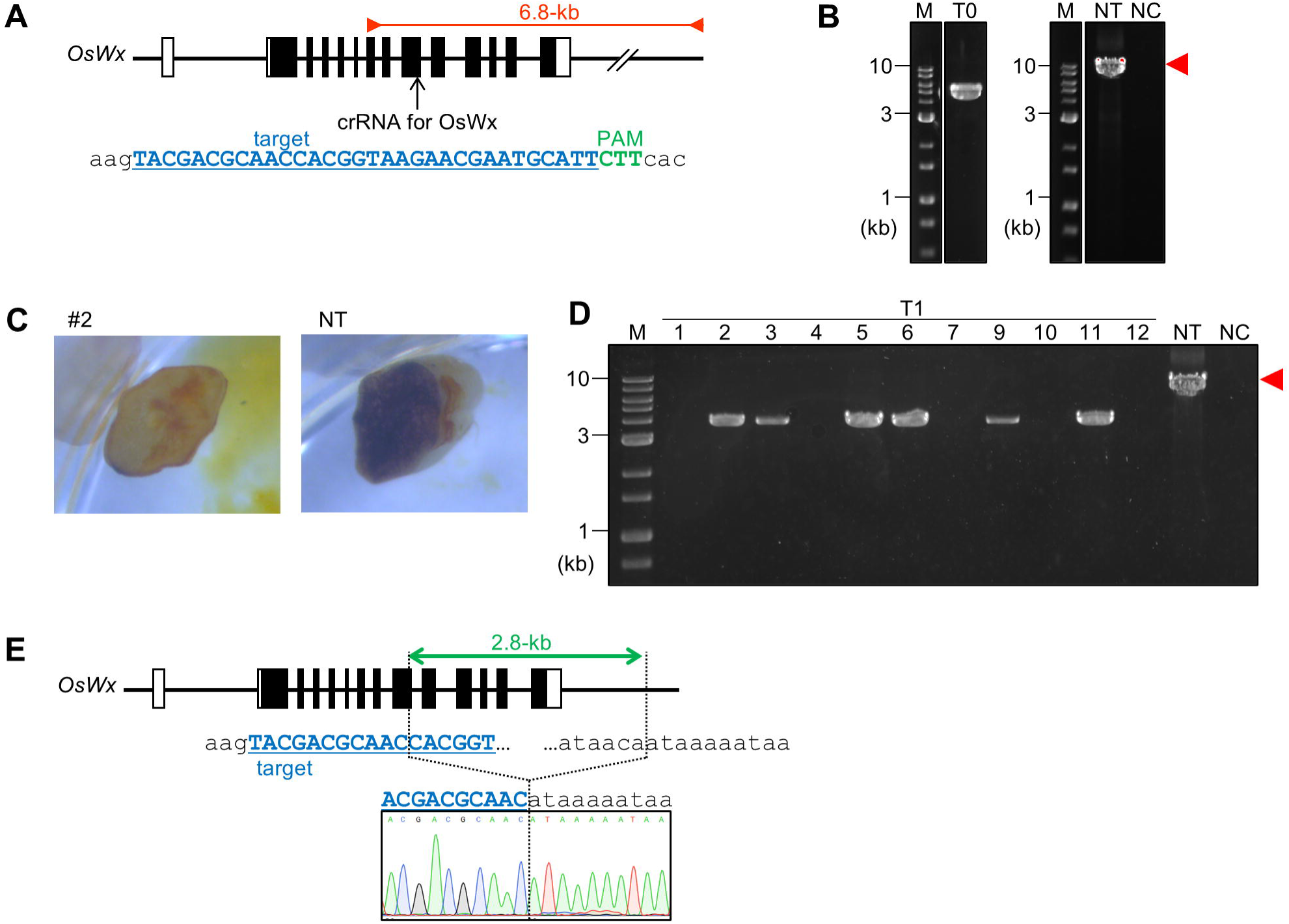
Inheritance in rice plants of mutations introduced by Eco CRISPR-Cas3. (A) Structure of *OsWx* locus. Details as in Fig. 1B. (B) PCR analysis of T0 plants transformed with Eco CRISPR-Cas3 for *OsWx*. Details as in Fig. 1D. Red arrowhead, fragment size amplified in wild type. (C) T1 seeds of #2 stained with iodine, NT; non-transformant. (D) PCR analysis of progenies in T1 generation. Red arrowhead, fragment size amplified in wild type. (E) Representative sequence chromatogram of #2 T1 plants.

Although non-transformed seeds were stained blue, none of the T1 seeds showed iodine staining, suggesting knockout of *OsWx* (Fig. 2C). PCR analysis in 11 T1 seedlings (except for #8, which did not germinate) showed that, compared with wild type, shorter fragments (6.8 kb) were amplified in 6 lines and no amplicons were amplified in 5 lines (Fig. 2D). Sanger sequencing of shorter amplicons confirmed a 2.8 kb deletion in the *OsWx* locus (Fig. 2E). These results showed that large deletions introduced by Eco CRISPR-Cas3 were inherited successfully to the next generation, as with TiD and Dvu I-C (Li et al. 2023; Osakabe et al. 2020). Considering that all T1 seeds showed waxy phenotype (Fig. 2C and D), it was assumed that biallelic mutations (2.8 kb deletion and no amplification) were introduced in T0 plants. Thus, it is estimated that biallelic mutants were obtained at 14% (1/7 regenerated plants) using Eco CRISPR-Cas3, suggesting that genome editing by Eco CRISPR-Cas3 is efficient enough to produce mutant rice plants.

### All-in-one vector for Eco CRISPR-Cas3

Here, 7 gene expression cassettes for 6 Cas effector proteins and crRNA of Eco CRISPR-Cas3 were prepared and carried separately on two binary vectors (Fig. 1A and Supplementary Fig. S1A). It was expected that all components could be expressed strongly using this strategy. However, the T-DNAs were large (15.7 kb and 14.2 kb in Cas537 and Cas6811+OsPDS, respectively) and both binary vectors need to be co-transformed into rice cells. This might represent a bottleneck to expanding Eco CRISPR-Cas3-mediated genome editing to plants in which it is harder to obtain transformed and genome-edited cells than in rice. In a previous report on induced pluripotent stem cells, an expression vector in which Eco CRISPR-Cas3 proteins were combined with 2A self-cleavage peptides was used to introduce deletions (Morisaka et al. 2019). In addition, vectors to express multiple Cas effector proteins of TiD and Dvu I-C combined with 2A peptides can be used for genome editing in plants (Li et al. 2023; Osakabe et al. 2020).

Thus, an all-in-one vector carrying 4 expression cassettes, Cas758 (EcoCas7, EcoCas5, and EcoCas8 combined with 2A peptides), Cas1163 (EcoCas11, EcoCas6, and EcoCas3 combined with 2A peptides), crRNA, and hygromycin phosphotransferase was constructed (Supplementary Fig. S3). Moreover, the crRNA expression cassette was inserted 400 bp away from the right border of T-DNA in the all-in-one vector, whereas it was inserted 90 bp from the right border of T-DNA in co-transformation vectors. An all-in-one vector harbors 15.9 kb of T-DNA. PCR analysis performed in rice calli transformed with all-in-one vectors targeting for *OsPDS* and *OsDL* estimated the frequency of transformed calli in which shorter amplicons were found at 55.3–70.8% (Supplementary Table S1).

Thus, in addition to co-transformation vectors carrying 7 expression cassettes of Eco CRISPR-Cas3, a compact all-in-one vector can also be applied to genome editing in rice. The deletion frequency seemed to be slightly higher compared with co-transformation with 2 binary vectors (Supplementary Table S1). This might be attributed to a combination of the following points. First, the crRNA expression cassette was protected in an all-in-one vector because regions adjacent to the border sequences in T-DNA are often lacking. Second, hygromycin and G418 were used for selection of cells co-transformed with Cas573 and Cas6811 vectors, whereas only hygromycin was used with the all-in-one vector. The number of cell divisions and cell activity are thought to differ between calli harboring these vectors, although the calli propagation period was the same (1.5 months). The propagation of hygromycin- and G418-resistant calli seemed to be slower than that of hygromycin-resistant calli. Speed of propagation might affect mutation frequency by Eco CRISPR-Cas3.

### Detailed analysis of mutation patterns induced by Eco CRISPR-Cas3

To characterize Eco CRISPR-Cas3-mediated deletions at the target site, the amplicons shown in Fig. 1D were cloned and sequenced. Sanger sequencing of clones selected randomly confirmed 8 types of mutations (a–h) (Fig. 3A). Deletions with a start point in the region between 278 bp upstream to 53 bp downstream of the PAM sequence occurred, and deletions were large (136–7218 bp). In previous reports on Type I-E CRISPR-Cas3 derived from *Thermobifida fusca* (Tfu CRISPR-Cas3) in human cells (Dolan et al. 2019) and Dvu I-C in maize (Li et al. 2023), mutation patterns were categorized into 4 groups: (Group I) one seamless junction, (Group II) one junction with insertion or partial inversion, (Group III) one junction with mutation(s), and (Group IV) two junctions. In this study, 3 and 4 types of mutations in Groups I and IV, respectively, were found. Types (a), (b), and (g) were single large deletions (6,617 bp, 7,218 bp, and 4,383 bp, respectively). On the other hand, mutations in types (c), (d), (e), and (f) consisted of 2 large deletions separated by intervening fragments of 8–361 bp. CRISPR-Cas3 induces DNA double-strand digestion by repetitive unwinding of target DNA and cleavage of DNA strands in *cis* and *trans* (Yoshimi et al. 2022). Cas3 can attack target sequences many times because, unlike Cas9, type I-E Cas3-mediated deletions often occur upstream of the PAM sequences (Dolan et al. 2019; Morisaka et al. 2019; Yoshimi and Mashimo 2022). This suggested the possibility that these mutations resulted from some rounds of EcoCas3-mediated deletion events. Surprisingly, type (h) consists of 4 large deletions separated by intervening fragments of 44, 103, and 124 bp that have not been found in previous reports (Dolan et al. 2019; Li et al. 2023). In types (d) and (e), a single large deletion concomitant with 85 bp and 4 bp of insertion, respectively, was observed. In type (e), inversion of a 156-bp fragment was observed. These unusual mutation patterns were also observed in Tfu CRISPR-Cas3-mediated genome editing in human cells (Dolan et al. 2019). Interestingly, a BLAST search showed that the 85 bp fragment in type (d) matched sequences located 23.8 kb upstream of the PAM sequence, suggesting the possibility that the genomic region including an 85 bp fragment 23.8 kb distant from PAM sequence accesses the target region on Eco CRISPR-Cas3 DNA digestion, although the detailed molecular mechanism is unclear. There were 2–6 bp microhomology sequences at the junctions in 7 of 15 patterns of deletion detected in this study, suggesting that microhomology-mediated end joining is involved in DNA double-strand repair in CRISPR-Cas3-mediated deletion as reported in Type I-C and I-D CRISPR-Cas3-mediated deletion in plants (Li et al. 2023; Osakabe et al. 2020).

**Figure 3.**
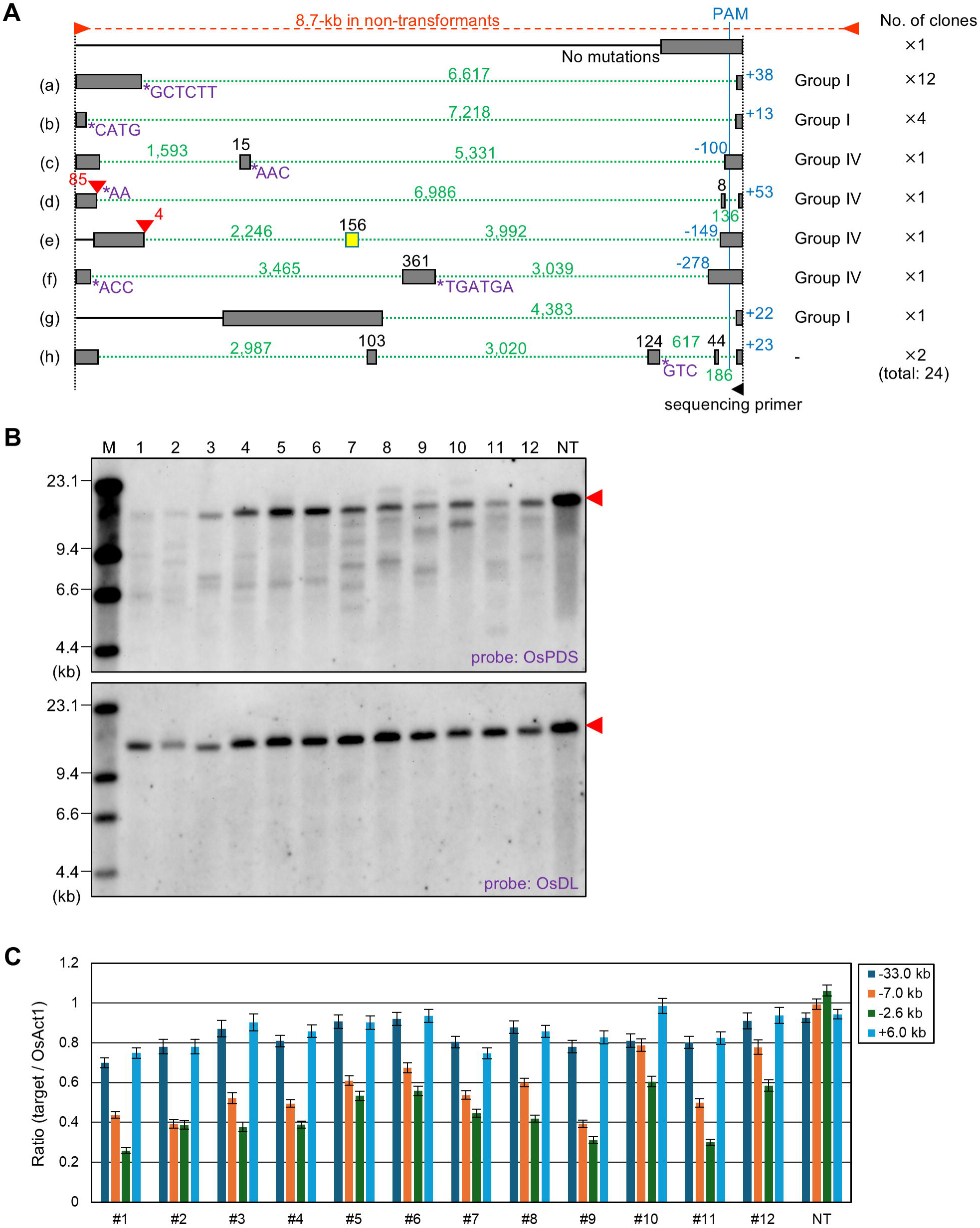
Deletions introduced by Eco CRISPR-Cas3 targeting *OsPDS* in rice calli. (A) Sequence analysis in transformed calli of #2. PCR fragments were sequenced with the primer shown (black arrowhead). Grey boxes and black solid lines indicate regions in which sequences were confirmed and not confirmed, respectively. A yellow box indicates an inverted region. Black numbers above grey and yellow boxes show their lengths. Green dotted lines/numbers and red arrowheads/numbers indicate regions/lengths of deletion and insertion, respectively. Blue numbers indicate the distance from the PAM at the site where the deletion started. Purple asterisks and letters indicate microhomology sequences at the junctions. (B) Southern blot analysis of *Sac*I-digested calli transformed Eco CRISPR-Cas3 with crRNAs for *OsPDS* using the probes shown in Fig. 1B and Fig. S1B. Arrowhead, band corresponding to wild type. M; size marker, NT; non-transformant. (C) Estimation of deletion size by ddPCR. The y-axis shows the ratio of amplicon concentrations at sites located 33.0 kb, 7.0 kb, and 2.6 kb upstream, and 6.0 kb downstream of the PAM sequence to those at the control site (*OsActin1*). Bars, technical error (maximum and minimum Poisson distribution for the 95% confidence interval).

It has been discussed that Dvu I-C tends to introduce “simple” large deletions compared with Eco CRISPR-Cas3 (Li et al. 2023). Also in the case of TiD, single large deletion and single large deletion with small insertion were determined in tomato and human cells, respectively, although bidirectional deletions were introduced, unlike type I-C and I-E (Osakabe et al. 2020; Osakabe et al. 2021). In this study, sequence analysis determined the presence of deletions concomitant with genome rearrangements such as short insertion and inversion in addition to simple deletions (Fig. 3A). This suggests that Eco CRISPR-Cas3 enables the production of both simple deletion and various kinds of alleles based on large deletions, although the frequency might be low. Rice calli stably transformed with Eco CRISPR-Cas3 were cultured for 1.5 months and analyzed in this study. On the other hand, CRISPR-Cas3 worked transiently in previous studies because expression vectors for Dvu I-C were transformed into maize protoplasts (Li et al. 2023) and ribonucleoprotein (RNP) of Tfu CRISPR-Cas3 infected human cells (Dolan et al. 2019). This suggests that continuous and repetitive Eco CRISPR-Cas3 attack on target regions in stably transformed calli results in longer and more complex mutation patterns. Optimization of callus culture conditions and period could potentially generate various mutation patterns. Promoter engineering by genome editing in agronomical important genes has been reported in crops such as tomato (Rodríguez-Leal et al. 2017) and rice (Cui et al. 2023). Dozens of gRNAs were prepared to create promoter deletion alleles in these reports. Eco CRISPR-Cas3 could also be used to create promoter alleles harboring large deletions with/without short insertion and inversion by use of a single gRNA in rice (Fig. 3A). Eco CRISPR-Cas3 could possibly induce more dynamic rearrangements because the deletion size and position are flexible compared with Class 2 CRISPR-Cas. In this scenario, technologies to introduce shorter deletions, i.e., dozens to hundreds of base pairs, might be needed to expand the application of Eco CRISPR-Cas3-mediated genome editing.

### Estimation of deletion frequency and length induced by Eco CRISPR-Cas3

Calli transformed with Eco CRISPR-Cas3 vectors are mosaics of various mutant cells. Electrophoresis of PCR products is a simple and easy way to find deletion events due to Eco CRISPR-Cas3. However, PCR bias makes it difficult to accurately calculate deletion frequencies or determine deletion length. Here, to estimate deletion frequency and length more precisely, the following analyses were performed. First, Southern blot analysis performed to visualize deletion events more clearly showed that shorter, and a few bands longer, than 14.1 kb corresponding to wild type were detected using a probe for *OsPDS* but not for *OsDL* as a control (Fig. 3B), supporting the view that deletions were introduced in this locus. Similar results were obtained in experiments using a different target designed in *OsDL* (Supplementary Fig. S1D). Second, the copy numbers of specific short regions around the target sequence were quantified using the droplet digital PCR (ddPCR) method, and the ratios to an internal control (*OsAct1*) were calculated (Fig. 3C). The ddPCR method was used to detect whether the target region was present or absent in experiments using Dvu I-C (Li et al. 2023). ddPCR can be used to estimate deletion length by quantification of presence or absence in multiple regions. Here, 3 regions at 33.0 kb, 7.0 kb, and 2.6 kb upstream of the PAM sequence and 1 region at 6.0 kb downstream of the PAM sequence were analyzed.

The ratios of copy number of a region 2.6 kb upstream of the PAM sequence to those in *OsAct1* were between 25.9% and 60.5% in all 12 samples analyzed, suggesting that the frequency of alleles lacking a region 2.6 kb upstream of the PAM sequence was 39.5–74.1%. Similarly, the frequency of alleles lacking a region at 7.0 kb upstream of the PAM sequence was 21.1–60.8%, although that frequency was 1.5– 25.2% in a region 6.0 kb downstream of the PAM sequence. This result is consistent with previous reports that type I-E CRISPR-Cas3 introduces uni-directional deletion in human cells (Dolan et al. 2019; Morisaka et al. 2019).

### Off-target mutations by Eco CRISPR-Cas3

The lower potential for off-target mutations due to the longer target sequence in Class 1 CRISPR-Cas compared to Class 2 CRISPR-Cas is considered one of the merits of Class 1 CRISPR-Cas (Yoshimi and Mashimo 2022). To investigate this, potential off-target sites of the 27-nt sequences used in this study were searched in the rice genome using the program GGGenome (https://gggenome.dbcls.jp/). This is because every 6th, 12th, 18th, 24th, and 30th base from the 5′ end of spacer sequences are not involved in target recognition, although the length of spacer sequence target is 32-nt in Eco CRISPR-Cas3. Sequences with mismatches of fewer than 6 nt (more than 20% mismatches) and potential PAM sequences (AAG, TAG, GAG, AGG, and ATG) were picked up (supplementary Table S2). As a result, 1 and 3 sites with 5-nt mismatches to spacer sequences in *OsPDS* and *OsDL* were identified, respectively. However, there are no potential PAM sequences at the 5′ end of spacer sequences. Moreover, 14, 36, and 1 sites with 6 nt mismatches to spacer sequences in *OsPDS*, *OsDL*, and *OsWx* were identified. Among them, 1 and 2 sites with 6-nt mismatches to spacer sequences harboring potential PAM sequences in *OsPDS* and *OsDL* were identified. PCR analysis was performed to confirm whether or not deletions were introduced in these off-target candidates. As expected, shorter bands than wild type were not detected, indicating that large deletions were not introduced by Eco CRISPR-Cas3 (Fig. 4 and Supplementary Fig. S4). Type I-E CRISPR-Cas has been reported to introduce no small indels (Dolan et al. 2019; Li et al. 2023; Morisaka et al. 2019; Yoshimi et al. 2024). Thus, this result suggested that off-target mutations were not introduced in these sites, although detailed analysis such as whole genome sequence may be needed.

**Figure 4.**
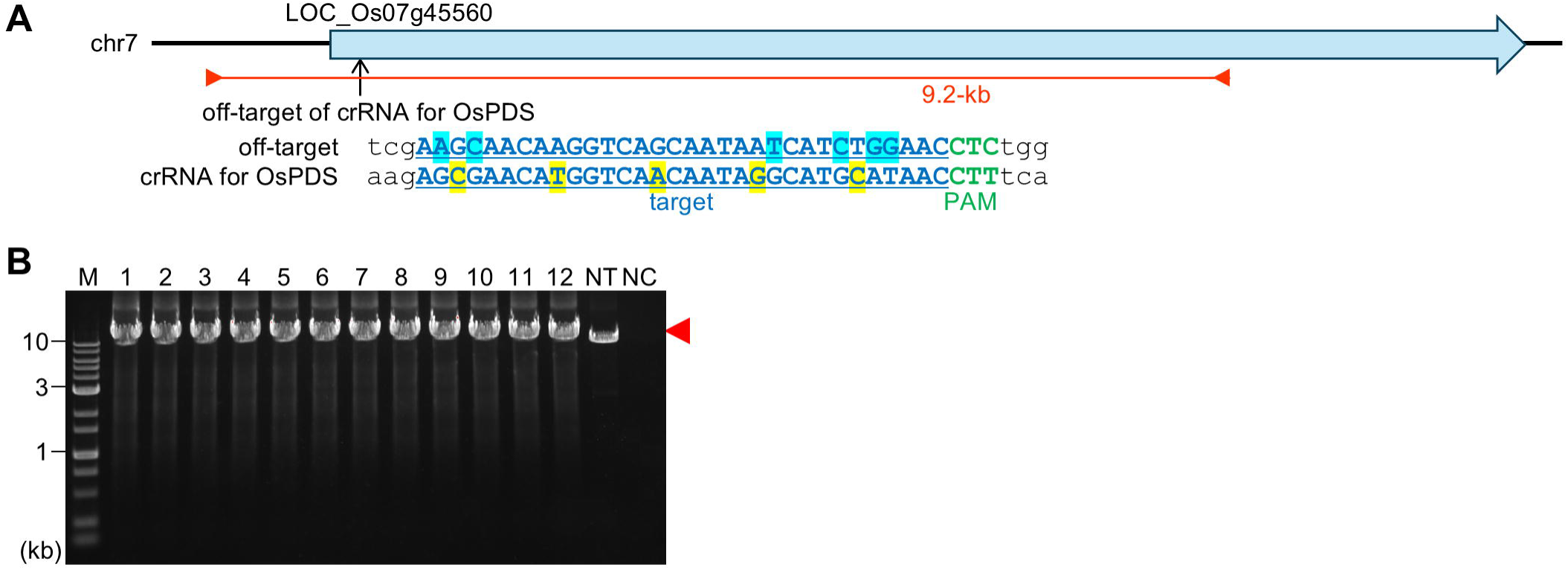
Off-target analysis of Eco CRISPR-Cas3-mediated targeted mutagenesis for *OsPDS* in rice calli. (A) Off-target locus of crRNA for *OsPDS* on chromosome 7. Mismatches and bases at 6, 12, 18, 24, and 30 nt from PAM that do not involve target recognition are highlighted in light blue and yellow, respectively. (B) PCR analysis of transformed calli using the primer set shown as orange arrowheads in A.

### Conclusion

In this study, we demonstrated targeted mutagenesis by Eco CRISPR-Cas3 in rice (Fig. 1 and Supplementary Fig. S1), and show inheritance of the introduced deletion to the progenies (Fig. 2 and Supplementary Fig. S2). Off-target mutations due to CRISPR-Cas3 can be kept at a low frequency because CRISPR-Cas3 recognizes longer target sequences compared with Class 2 CRISPR-Cas. Off-target mutations, large deletions in this case, were not detected in rice (Fig. 4). As reported previously in human cells (Dolan et al. 2019; Morisaka et al. 2019), several kb of uni-directional deletion can be introduced into the target region by Eco CRISPR-Cas3 in rice (Fig. 3A and C). Moreover, deletions separated by intervening fragments and deletions with short insertion and inversion were also determined (Fig. 3A). Our results suggest that targeted mutagenesis using Eco CRISPR-Cas3 can create novel gene alleles in addition to gene knockouts and deletions.

## Materials and Methods

### Oligonucleotides

Primers used in this study are listed in Supplementary Table S3.

### Vector construction

#### Co-transformation vectors

To construct vectors for Eco CRISPR-Cas expression cassettes, rice codon optimized EcoCas3, EcoCas5, EcoCas6, EcoCas7, EcoCas8, and EcoCas11-coding sequences including bpNLS were synthesized by GeneArt Gene Synthesis (Thermo Fisher Scientific). These vectors inserted into *Spe*I/*Sac*I-digested pE(R4-R3) ZmUbi_OsFnCpfI_Tpea3A (Endo et al. 2016) by an In-fusion reaction (Takara), yielding vectors for Cas expression driven by the maize ubiquitin promoter. PCR fragment amplified fragments (using primer sets, attB1 f/attB4 r, attP4 f/attP3 r, and attB3 f/attB2 r) with these vectors as templates were cloned into entry clones, pDONR221 P1-P4, pDONR221 P4r-P3r, and pDONR221 P13-P2, respectively, by BP reaction (Invitrogen), yielding pDONR221(L1-L4)ZmUbiCas5Tpea3A, pDONR221(L1-L4)ZmUbiCas6Tpea3A, pDONR221(R4-R3)ZmUbiCas3Tpea3A, pDONR221(R4-R3)ZmUbiCas8Tpea3A, pDONR221(L3-L2)ZmUbiCas7Tpea3A, and pDONR221(L3-L2)ZmUbiCas11Tpea3A (Supplementary Sequence). To construct a destination vector harboring an nptII-expression cassette, an nptII fragment amplified with primer set nptII f2/nptII r2 was inserted into *Xho*I-digested pZD202Hyg vector (Kwon et al. 2012) by an In-fusion reaction, yielding pZD202Km. Entry clones were cloned into pZD202Hyg and pZD202Km using LR clonase II plus, yielding Cas537 and Cas6811 (pZD202Hyg harboring Cas5, Cas3, and Cas7 expression cassettes and pZD202Km harboring Cas6, Cas8, and Cas11 expression cassettes, respectively). To construct a crRNA expression vector, a pOsU6Cas3 gRNA vector harboring OsU6-2 promoter, 86-nt leader sequence, and two 29-nt repeat sequences were synthesized by GeneArt Gene Synthesis. Annealed oligonucleotides in Supplementary Table S3 were ligated into *Bsa*I-digested pOsU6Cas3 gRNA (Supplementary Sequence). The crRNA expression cassettes amplified with a primer set, gRNA PmeI f/gRNA PmeI r were inserted into *Pme*I-digested Cas537 and Cas6811, yielding Cas537+OsDL, Cas6811+OsPDS, and Cas6811+OsWx.

### All-in-one vector

A pE(L3-L2)P2X35S::I-SceI::Thsp vector (Kwon et al. 2012) was digested with *Asc*I and *Pac*I, and inserted into *Asc*I-/*Pac*I-digested pDONR221(L1-L4) vector, yielding pDONR221(L1-L4)2×35S-I-SceIos-T17.3. Four PCR fragments amplified using primer sets P35S-omega f1/RcCAT r2, bpNLS f1/T2A-NLS r1, T2A-NLS f1/T2A-NLS r2, T2A-NLS f2/T17.3 r1 with pDONR221(L1-L4)2×35S-I-SceIos-T17.3, pDONR221(L3-L2)ZmUbiCas7Tpea3A, pDONR221(L1-L4)ZmUbiCas5Tpea3A, and pDONR221(R4-R3)ZmUbiCas8Tpea3A, respectively, as templates were cloned into *Xba*I-/*Sac*I-digested pDONR221(L1-L4)2×35S-I-SceIos-T17.3 by an In-fusion reaction, yielding pDONR221(L1-L4) 2×35SΩint Cas758 T17.3 (Supplementary Sequence). Three PCR fragments amplified using primer sets ADHUTR-NLS f1/T2A-NLS r3, T2A-NLS f2/r1, T2A-NLS f1/Tpea3A r5 with pDONR221(L3-L2)ZmUbiCas11Tpea3A, pDONR221(L1-L4)ZmUbiCas6Tpea3A, and pDONR221(R4-R3)ZmUbiCas3Tpea3A, respectively, as templates were cloned into *Spe*I-/*Sac*I-digested pDONR221(R4-R3)ZmUbiCas8Tpea3A by an In-fusion reaction, yielding pDONR221(R4-R3)ZmUbiCas1163Tpea3A (Supplementary Sequence). Three entry clones, pDONR221(L1-L4) 2×35SΩint Cas758 T17.3, pDONR221(R4-R3)ZmUbiCas1163Tpea3A, and pDONR221(L3-L2)NoshptT35Snos harboring hpt driven by the Nos promoter and 35S/Nos double terminator were cloned into pZD202 (Kwon et al. 2012) using a LR clonase II plus, yielding pZD202 Cas758+Cas1163+hpt. The crRNA expression cassettes amplified with a primer set gRNA SpeI f/gRNA SpeI r were inserted into *Spe*I-digested pZD202 Cas758+Cas1163+hpt, yielding all-in-one+OsDL, all-in-one+OsPDS.

### Transformation

For rice transformation, binary vectors were transformed into *Agrobacterium tumefaciens* strain EHA105 (Hood et al. 1993) by the electroporation method. *Agrobacterium*-mediated transformation of rice (*Oryza sativa* L. cv. Nipponbare) followed our previous reports (Toki 1997; Toki et al. 2006). Briefly, 4-week-old secondary calli derived from mature seeds were co-cultivated with *Agrobacterium* (mixed *Agrobacterium* harboring a different binary vector in case of co-transformation) for 3 days. Calli were washed to eliminate *Agrobacterium* and cultured on an N6D selection medium containing 50 mg/L hygromycin, 35 mg/L G418, and 25 mg/L meropenem in the case of co-transformation, or 50 mg/L hygromycin, and 25 mg/L meropenem in the case of an all-in-one vector for 6 weeks. Calli were transferred to fresh selection medium every 2 weeks. Calli growing vigorously on the selection medium were cultured on regeneration medium (ReIII) containing 25 mg/L meropenem, and shoots arising from calli were transferred to hormone-free medium containing 25 mg/L meropenem.

### DNA extraction and PCR analysis

Genomic DNA was extracted from transformed calli and leaves of rice using Agencourt chloropure (Beckman Coulter) or Nucleon Phytopure extraction kit (Cytiva) according to the manufacturer’s protocol. PCR analysis was performed with KOD One PCR master mix (TOYOBO) using the primer sets listed in Supplementary Table S3.

### Southern blot analysis

Southern blot analysis was performed by following a conventional protocol. *Sac*I-digested genomic DNA (5 μg) from calli was electrophoresed on a 0.7% gel at around 50 V and transferred to positively charged nylon membranes (Roche Diagnostics). Specific DNA probes were prepared using a PCR digoxigenin (DIG) probe synthesis kit (Roche Diagnostics) according to the manufacturer’s protocol using the primer sets listed in Supplementary Table S3. The probe-hybridized membranes were washed using DIG Wash and Block Buffer Set (Roche Diagnostics). Chemical luminescence on the membrane with CDP-Star treatment was detected with a ChemiDoc Touch (BIO-RAD).

### ddPCR

For ddPCR, 22 µl of reaction mixture containing 2.5 ng of genomic DNA digested with *Pac*I and *Sac*I, ddPCR Supermix for Probes (no dUTP) (Bio-Rad), and PrimePCR Probe Assay (Bio-Rad) were prepared. In this study, PrimePCR Probe Assay: ACT1, Rice labeled with HEX, PrimePCR Probe Assay: OS03G0183500 *, Rice labeled with FAM (–33.0 kb), PrimePCR Probe Assay: OS03G0183900 *, Rice labeled with FAM (–7.0 kb), PrimePCR Probe Assay: PDS, Rice labeled with FAM (–2.6 kb), and PrimePCR Probe Assay: OS03G0184100 *, Rice labeled with FAM (+6.0kb) were used (Bio-Rad). Droplet generation using an Automated Droplet Generator (Bio-Rad), PCR using droplets, droplet analysis using a QX200 Droplet Reader (Bio-Rad), and data analysis with QuantaSoft software (Bio-Rad) followed our previous report (Nishizawa-Yokoi et al. 2021).

### Sequencing analysis

For Sanger sequence analysis of PCR fragments, PCR products were cloned into pCR-Blunt II-TOPO (Invitrogen) and transformed into *E. coli*. Colony PCR was performed with KOD One PCR master mix (TOYOBO) using M13 forward and reverse primers. The resultant PCR products were used as a template for sequencing reaction using a BigDye Terminator v3.1 Cycle Sequencing Kit (Thermofisher Scientific) and subjected to sequence analysis using an ABI3130 sequencer (Thermofisher Scientific). Sequence data were analyzed using SnapGene (GSL Biotech LLC).

## Data Availability Statement

The data supporting this study are available in the article and supplementary data.

## Acknowledgments

We thank Drs. M. Endo, A. Nishizawa-Yokoi, and S. Hirose for critical discussion and Dr. Helen Rothnie for English editing.

## Author contributions

H.S. and S.Y. designed the experiments. H.S., H.N. conducted the experiments and analyzed the data. T. Muranaka, K.Y., T. Mashimo, and S.T. supervised the research. H.S. wrote the manuscript. All authors read and approved of the manuscript.

