## Supplementai File for "Heritable Large Deletions Using Type I-E CRISPR-Cas3 in Rice"

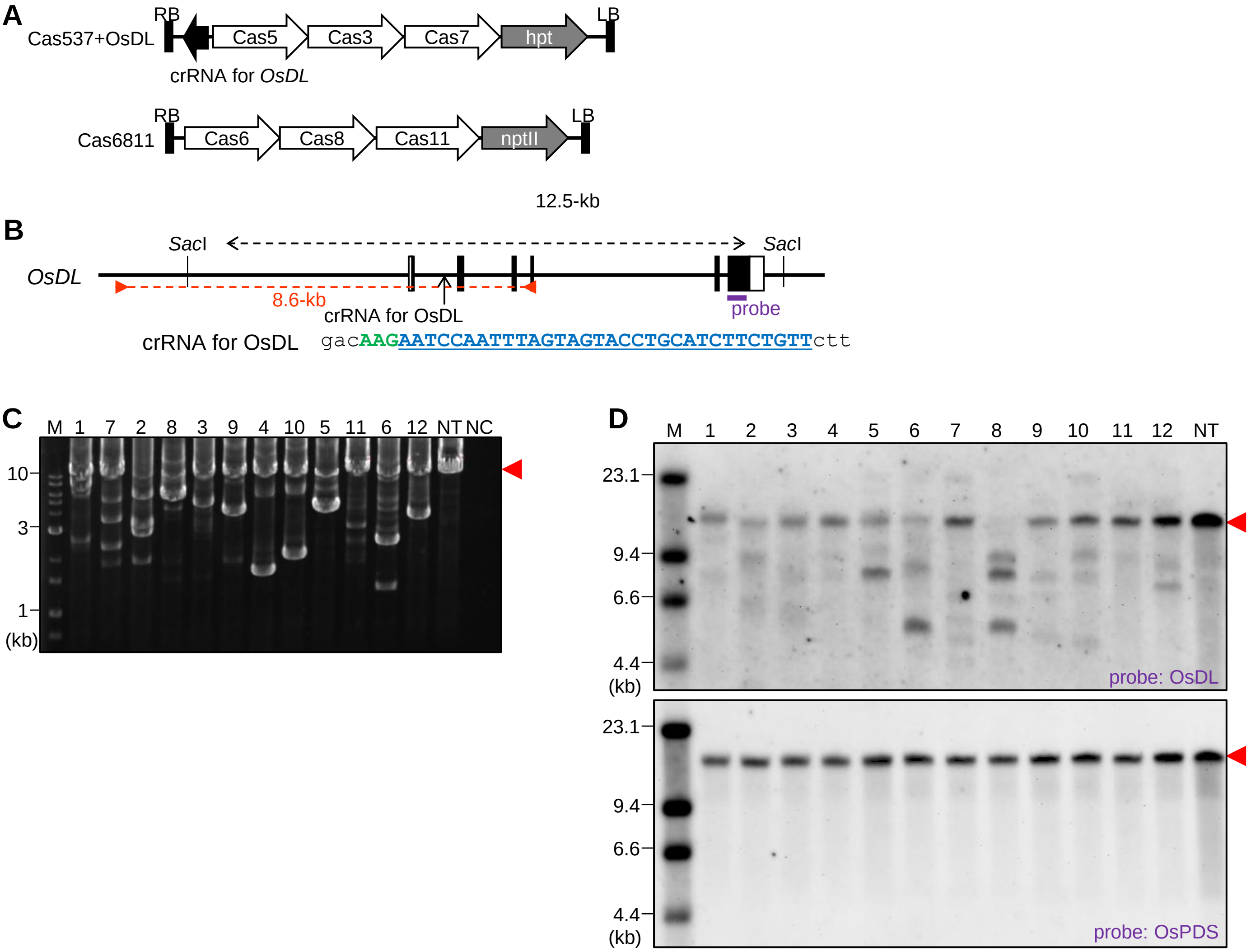


**Supplementary Figure S1** Eco CRISPR-Cas3-mediated targeted mutagenesis for *OsDL* in rice calli. (A) Eco CRISPR-Cas3 vectors targeting for *OsDL*. (B) Structure of *OsDL* locus. (C and D) PCR and Southern blot analysis of transformed calli. Details are same as Fig. 1A, B, D and 3B.

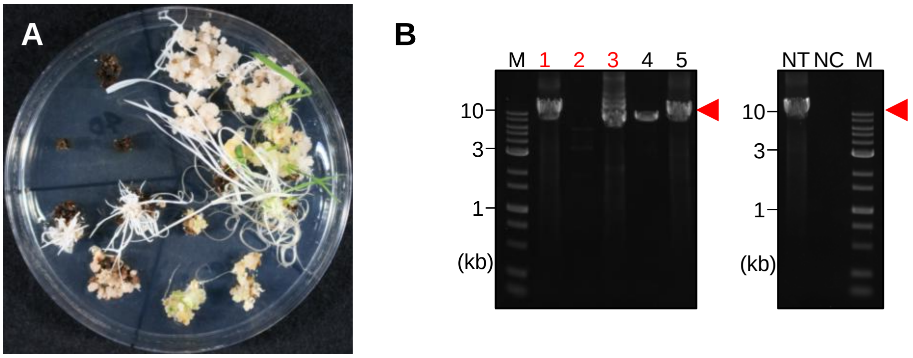


**Supplementary Figure S2** Eco CRISPR-Cas3-mediated targeted mutagenesis for *OsPDS* in regenerated plants.
(A) Rice regenerated plants transformed with binary vectors harboring Eco CRISPR-Cas3 with crRNAs for *OsPDS* shown in Fig. 1A. (B) PCR analysis in regenerated plants. Red numbers indicate albino lines. Details are same as Fig. 1D.

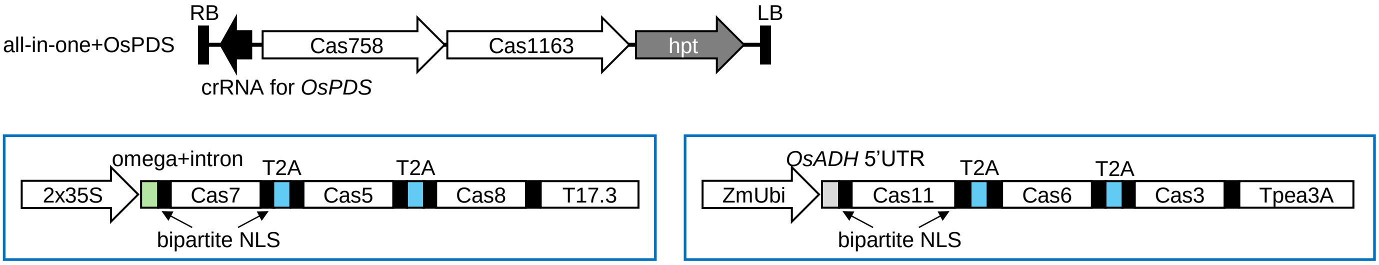


**Supplementary Figure S3** An all-in-one vector of Eco CRISPR-Cas3 targeting for *OsPDS*
The detail construction of Cas758 and Cas1163 expression cassettes are shown in boxes with blue lines. A green box shows CaMV omega sequence and caster bean catalase intron. Blue boxes shows T2A peptides. 2x35S; double CaMV 35S promoter, T17.3; rice heat shock protein 17.3 terminator. Details are same as Fig. 1A.

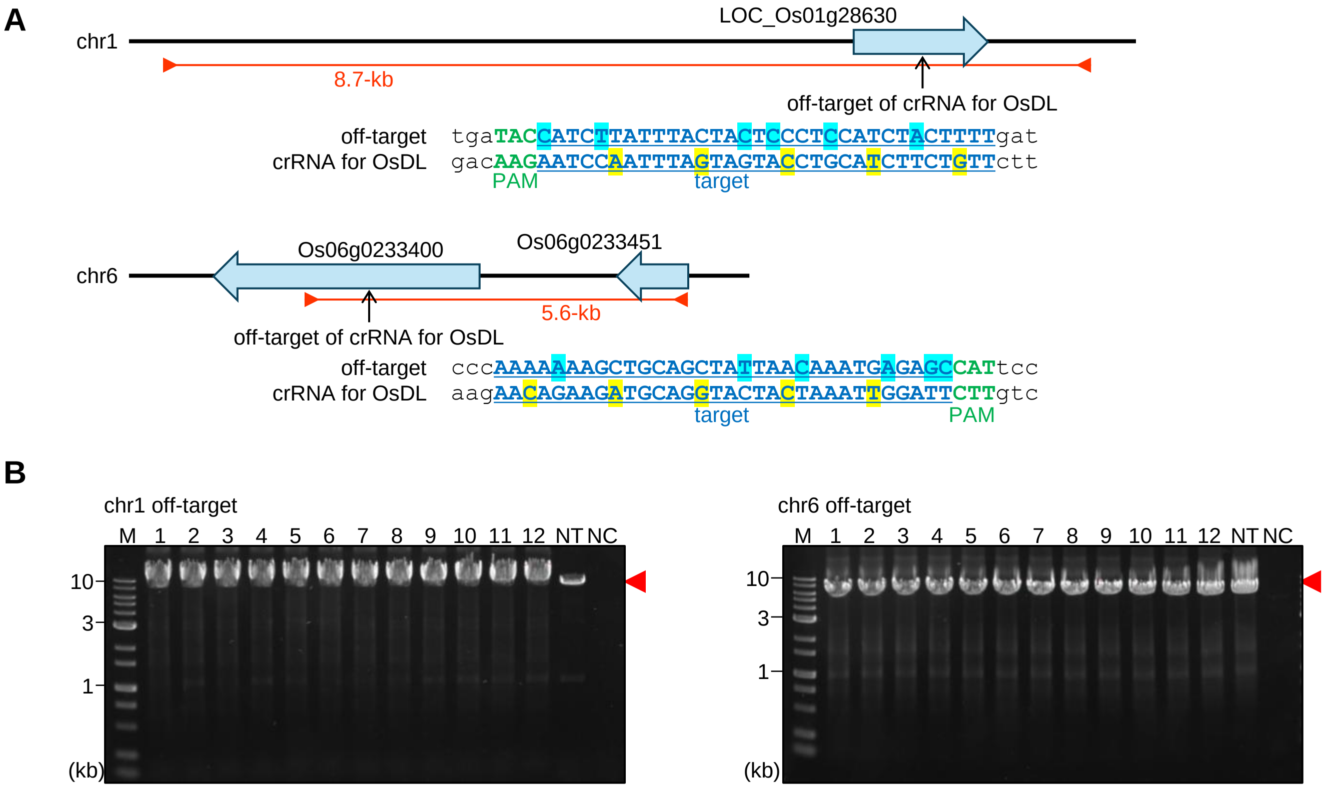


**Supplementary Figure S4** Off-target analysis of Eco CRISPR-Cas3-mediated targeted mutagenesis for *OsDL* in rice calli.
(A) Off-target locus of crRNA for *OsDL* on chromosome 1 and 6. (B) PCR analysis of transformed calli using a primer set shown in A.

**Supplementary Table S1**. Targeted mutagenesis frequency using an all-in-one vector.

| target | vector | No. of calli in which shorter fragments were found (A) | No. of calli in which fragments were amplified (B) | mutation frequency (A/B) |
|---|---|---|---|---|
| OsPDS | Cas537/Cas6811+OsPDS | 13 | 33 | 39.4% |
| | | 21 | 44 | 47.7% |
| | | 27 | 56 | 48.2% |
| | all-in-one+OsPDS | 31 | 45 | 68.9% |
| | | 34 | 48 | 70.8% |
| OsDL | Cas537+OsDL/Cas6811 | 10 | 32 | 31.3% |
| | | 12 | 42 | 28.6% |
| | all-in-one+OsDL | 38 | 54 | 70.4% |
| | | 26 | 47 | 55.3% |

**Supplementary Table S2**. List of potential off-target sites.

(A) Summary

| Target | query | No. of 5-nt mismatch (with potential PAM) | No. of 6-nt mismatch (with potential PAM) | No. of 7-nt mismatch (with potential PAM) |
|---|---|---|---|---|
| OsPDS | GTTATnCATGCnTATTGnTGACCnTGTTCnCT | 5nt-mismatch: 1 (0) | 13 (1) | 154 (11) |
| OsDL | AATCCnATTTAnTAGTAnCTGCAnCTTCTnTT | 5nt-mismatch: 3 (0) | 34 (2) | 402 (60) |
| OsWx | AATGCnTTCGTnCTTACnGTGGTnGCGTCnTA | 0 (0) | 1 (0) | 36 (2) |

(B) OsPDS (6-nt and 7-nt mismatches with potential PAM)

| chromosome | strand | start | sequence | Mismatch sites (X) | No. of mismatch |
|---|---|---|---|---|---|
| chr02 | + | 7300166 | GTTGTTTTTTCTTATTCTTGACCATGATCTCC | ===X==XX=X======X=========X====X | 7 |
| chr02 | + | 8113462 | GCTTTTCATGCATATTGCTTCCTCTCTTTGCT | =X=X===============XX=X==X==X=== | 7 |
| chr05 | + | 25696094 | GTTTTTCATGCATGATGATAACCATGCACGCA | ===X=========XX====X======XX===X | 7 |
| chr07 | + | 4077425 | GCCATCCAGGCATAATGTTGACCATGCACGGT | =XX=====X=====X===========XX==X= | 7 |
| chr08 | + | 5466569 | GTTCTCCATGCCTTTTGGTGAATCAGCTAGCT | ===X=========X=======XX=X=X=X=== | 7 |
| chr08 | + | 17689275 | GTTATACATCATCTATGGAGTCCATGTTCACT | =========XX=XXX===X=X=========== | 7 |
| chr08 | + | 22279644 | CTGATACACGCTTGTTGTTGTTGTTGTTCCCT | X=X=====X====X======XXX========= | 7 |
| chr11 | + | 7955664 | GGTATTCATGTATAATTTTAACGGTGTTATCT | =X========X===X=X==X==X=====X=== | 7 |
| chr02 | - | 215829 | AGATAACAAGCTCTCTAATATATATGAGTAAC | ===X======X==X=X=====XX====X==== | 7 |
| chr05 | - | 9825229 | ACTCAACAAGGTCACCATCAAGAATAAATATC | =X=X============XX===X==X====X= | 7 |
| chr07 | - | 27179127 | AAGCAACAAGGTCAGCAATAATCATCTGGAAC | =X=X================X===X=XX=== | 6 |
| chr09 | - | 21854990 | AGATAACACCGTTAGAATTAGACATGAATACC | ===X=====X==X==X=X===X========X= | 7 |

(C) OsDL (6-nt and 7-nt mismatches with potential PAM)

| chromosome | strand | start | sequence | Mismatch sites (X) | No. of mismatch |
|---|---|---|---|---|---|
| chr01 | + | 2034952 | AATACAATTTATAAATAACTAAAACTTGTGAT | ===X========X=X=====XX=====X==X= | 7 |
| chr01 | + | 2470183 | AATACAATTTAGAAGTAACTGAAATTTGAAAT | ===X========X========X==X==XX=X= | 7 |
| chr01 | + | 15196524 | AATACAATTTAGAAGTAACTGAAATTTAAAAT | ===X========X========X==X==XX=X= | 7 |
| chr01 | + | 16020905 | CATCTTATTTACTACTCCCTCCATCTACTTTT | X===X=========X=X===X=====X===== | 6 |
| chr01 | + | 20158194 | CATGCAATTAACTAGCACATCCCTCTTCTTTT | X==X=====X=====X==X=X=X========= | 7 |
| chr01 | + | 22630010 | AATACAATTTAGAAGTAACTGAAATTTAAAAT | ===X========X========X==X==XX=X= | 7 |
| chr01 | + | 26204479 | AATACAATTTAGAAGTAACTGAAATTTGAAAT | ===X========X========X==X==XX=X= | 7 |
| chr01 | + | 26205077 | AATACAATTTAGAAGTAACTGAAATTTGAAAT | ===X========X========X==X==XX=X= | 7 |
| chr01 | + | 28528811 | AATACAATTTAGAAGTAACTGAAAATTCGAAA | ===X========X========X==X===X=XX | 7 |
| chr01 | + | 34197201 | AATACAATTTAGAAGTAACTGAAATTTGAAAT | ===X========X========X==X==XX=X= | 7 |
| chr01 | + | 34197804 | AATACAATTTAGAAGTAACTGAAATTTGAAAT | ===X========X========X==X==XX=X= | 7 |
| chr02 | + | 17269379 | AATACAATTTAGAAGTAACTGAAATTTGAAAT | ===X========X========X==X==XX=X= | 7 |
| chr02 | + | 21448444 | AATACAATTTAGAAGTAACTGAAATTTGAAAT | ===X========X========X==X==XX=X= | 7 |
| chr02 | + | 31168273 | AATACAATTTAGAAGTAACTGAAATTTGAAAT | ===X========X========X==X==XX=X= | 7 |
| chr02 | + | 33688077 | AATACAATTTAGAAGTAACTGAAATTTGAAAT | ===X========X========X==X==XX=X= | 7 |
| chr03 | + | 14882720 | ATTACCATTCAGATGTAACTACAGCTTCAGTT | =X=X=====X==XX======X=======X=== | 7 |
| chr03 | + | 15438619 | AATATGATTCACTAGTTTCTGCAGATTCAGTG | ===XX====X======X=======X===X==X | 7 |
| chr03 | + | 31516263 | AATACAATTTAGAAGTAACTGAAATTTGAAAT | ===X========X========X==X==XX=X= | 7 |
| chr04 | + | 543441 | AATACAATTTATAAATAACTGAAATTTGTGAT | ===X========X=X======X==X==X==X= | 7 |
| chr04 | + | 669321 | AATACAATTTAGAAGTAACTGAAATTTAAAAT | ===X========X========X==X==XX=X= | 7 |
| chr04 | + | 16503467 | AATACAATTTAGAAGTAACTGAAATTTGAAAT | ===X========X========X==X==XX=X= | 7 |
| chr04 | + | 25452654 | AATACAATTTAGAAGTAACTGAAATTTGAAAT | ===X========X========X==X==XX=X= | 7 |
| chr04 | + | 27141187 | AATACAATTTATAAGTAACTCAATTTTTTGAT | ===X========X=======XX==X==X==X= | 7 |
| chr04 | + | 30370638 | AATACAATTTAGAAGTAACTGAAATTTGAAAT | ===X========X========X==X==XX=X= | 7 |
| chr04 | + | 32978125 | AGTCCTTTTTGGAAGGAGCGGGAGCTTCTATT | =X====X===X=X==X===X=X========== | 7 |
| chr04 | + | 35110737 | AATACAATTTAGAAGTAACTGAAATTTGAAAT | ===X========X========X==X==XX=X= | 7 |
| chr05 | + | 19824739 | AATACAATTTATTAATAACTAAAATTTTTGAT | ===X==========X=====XX==X==X==X= | 7 |
| chr05 | + | 20054482 | AATACAATTTAGAAGTAACTGAAATTTAAAAT | ===X========X========X==X==XX=X= | 7 |
| chr05 | + | 22525707 | AATACAATTTAGAAGTAACTGAAATTTGAAAT | ===X========X========X==X==XX=X= | 7 |
| chr05 | + | 25479095 | AATACAATTTATAAATAACTAAAATTTGTGTT | ===X========X=X=====XX==X==X==== | 7 |
| chr05 | + | 26589223 | CATGCTATTAACTAGTTTCTCATTCTTCTTTT | X==X=====X======X===XXX========= | 7 |
| chr05 | + | 28719717 | AATACAATTTAGAAGTAACTGAAATTTGAAAT | ===X========X========X==X==XX=X= | 7 |
| chr06 | + | 429985 | AGTTGGATTTTTAAGTAGTTACAGCTTCTGTT | =X=XX=====X=X=====X=X========== | 7 |
| chr06 | + | 3664979 | AATACAATTTATAAATAACTAAAACTTGTGAT | ===X========X=X=====XX=====X==X= | 7 |
| chr07 | + | 81360 | AAGCGAATGGATTATTGACTGTATCTTCTTTT | ==X=X===XX====X=X====X========== | 7 |
| chr07 | + | 448581 | TAACATCTTTTCTAGTATCTGCATCTACATTT | X=X=X=X===X===============X=X=== | 7 |
| chr07 | + | 22226504 | AATACAATTTAGAAGTAACTGAAATTTGAAAT | ===X========X========X==X==XX=X= | 7 |
| chr07 | + | 29205777 | AATACAATTTAGAAGTAACTGAAATTTGAAAT | ===X========X========X==X==XX=X= | 7 |

(C) OsDL (continued)

| chromosome | strand | start | sequence | Mismatch sites (X) | No. of mismatch |
|---|---|---|---|---|---|
| chr08 | + | 7301278 | AATACAATTTAGAAGTAACTGAAATTTGAAAT | ===X========X========X==X==XX=X= | 7 |
| chr08 | + | 19671972 | AAGCCATTTTAGTACTCCCTCCATCTATTTTT | ==X===X=======X=X===X=====XX==== | 7 |
| chr08 | + | 26639861 | AATACAATTTAGAAGTAACTGAAATTTGAAAT | ===X========X========X==X==XX=X= | 7 |
| chr09 | + | 16579704 | ACTTTTATTTACTACTCCCTCCATCTACTTTT | =X=XX=========X=X===X=====X===== | 7 |
| chr09 | + | 22429519 | AATACAATTTAGAAGTAACTGAAATTTAAAAT | ===X========X========X==X==XX=X= | 7 |
| chr10 | + | 2002123 | AATACAATTTAGAAGTAACTGAAATTTGAAAT | ===X========X========X==X==XX=X= | 7 |
| chr11 | + | 129508 | AATACAATTTAGAAGTAACTGAAATTTGAAAT | ===X========X========X==X==XX=X= | 7 |
| chr11 | + | 7002819 | AATACAATTTAGAAGTAACTGAAATTTGAAAT | ===X========X========X==X==XX=X= | 7 |
| chr11 | + | 7003960 | AATACAATTTAGAAGTAACTGAAATTTGAAAT | ===X========X========X==X==XX=X= | 7 |
| chr11 | + | 23681933 | AATACAATTTAGAAGTAACTGAAATTTGAAAT | ===X========X========X==X==XX=X= | 7 |
| chr11 | + | 26275293 | AATACAATTTAGAAGTAACTGAAATTTGAAAT | ===X========X========X==X==XX=X= | 7 |
| chr11 | + | 26286307 | AATACAATTTAGAAGTAACTGAAATTTGAAAT | ===X========X========X==X==XX=X= | 7 |
| chr11 | + | 26956516 | AATACAATTTAGAAGTAACTGAAATTTGAAAT | ===X========X========X==X==XX=X= | 7 |
| chr11 | + | 27810188 | AATACAATTTAGAAGTAACTGAAATTTGAAAT | ===X========X========X==X==XX=X= | 7 |
| chr12 | + | 22300678 | AATAAAATTTAGTAGTAACTGAAATTTGAAAT | ===XX================X==X==XX=X= | 7 |
| chr01 | - | 30662713 | ATTTGAAATTTCAGTTATTTGTAAATTGTATT | =X=X===X==X======X=X========X=== | 7 |
| chr01 | - | 30709740 | ATTTCAAATTTCAGTTACTTCTAAATTGTATT | =X=XX==X==X========X========X=== | 7 |
| chr01 | - | 30710512 | ATTTCAAATTTCAGTTACTTCTAAATTGTATT | =X=XX==X==X========X========X=== | 7 |
| chr01 | - | 38095368 | ATTTCAAATTTCAGTTACTTCTAAATTGTATT | =X=XX==X==X========X========X=== | 7 |
| chr01 | - | 39519445 | ATTTCAAATTTCAGTTACTTCTAAATTGTATT | =X=XX==X==X========X========X=== | 7 |
| chr01 | - | 42907999 | ATTTCAAATTTCAGTTACTTCTAAATTGTATT | =X=XX==X==X========X========X=== | 7 |
| chr02 | - | 181460 | ATTTCGAATTTCAGTTACTAATAAATTGTATT | =X=XXX=X==X=================X=== | 7 |
| chr02 | - | 297312 | ATTTCGAATTTCAGTTACTAATAAATTGTATT | =X=XXX=X==X=================X=== | 7 |
| chr02 | - | 401935 | ATTTCGAATTTCAGTTACTAATAAATTGTATT | =X=XXX=X==X=================X=== | 7 |
| chr02 | - | 4318166 | ATTTCAAATTTCAGTTACTTCTAAATTGTATT | =X=XX==X==X========X========X=== | 7 |
| chr02 | - | 8270648 | ATTTCAAATTTCAGTTACTTCTAAATTGTATT | =X=XX==X==X========X========X=== | 7 |
| chr02 | - | 9286929 | ATTTTAAATTTCAGTTACTTCTAAATTGTATT | =X=XX==X==X========X========X=== | 7 |
| chr02 | - | 17092951 | AATTTTAATTTCAGTTACTTCTAAATTGTATT | ===XXX=X==X========X========X=== | 7 |
| chr02 | - | 18876065 | AATCTAAATTGAAAATATTAATAAATTGAATT | ===XX==X===X=X===X==========X=== | 7 |
| chr02 | - | 20469999 | AATTTAACAATGAGTTACTACTAAATGCGATT | ===XX==X=XXX===============X==== | 7 |
| chr02 | - | 20601835 | AAAAATAGATGGAGGGAATACTATATTGCATT | ====XX=====X===X=X=====X====X=== | 7 |
| chr02 | - | 21465036 | ATTACGAATTTCAGTTACTTCTAAATTGTATT | =X==XX=X==X========X========X=== | 7 |
| chr02 | - | 22088483 | AAAAGTAGATGGAGGAAGTAGTACATTATATT | =====X=====X===X=X=====X===XX=== | 7 |
| chr02 | - | 24660138 | ATTCCAAATTTCAGCTATTATTAAATTGTATT | =X=XX==X==X======X==========X=== | 7 |
| chr02 | - | 30360952 | ATTTCAAATTTCAGTTACTTCTAAATTGTATT | =X=XX==X==X========X========X=== | 7 |
| chr02 | - | 31066147 | ACCAGGAGCGGCAGGAAGAAGTAAATCGGCTT | =X===X===X=====X=XX==========X== | 7 |
| chr03 | - | 9888185 | ATCACAAGTTTTAGTTATTTATAAATTGTATT | =X==X=====XX=====X=X========X=== | 7 |
| chr03 | - | 16759604 | TAGAGAAGATGAAGATGCTAGTTAACTGAATG | X==========X====X=====X==X==X==X | 7 |
| chr03 | - | 23063891 | ATTTCAAATTTCAGTTACTTTTAAATTGTATT | =X=XX==X==X========X========X=== | 7 |
| chr03 | - | 25975657 | ATTTTAAATTTCAGTTACTTCTAAATTGTATT | =X=XX==X==X========X========X=== | 7 |
| chr03 | - | 29034316 | ATTTCAAATTTCAGTTACTTCTAAATTGTATT | =X=XX==X==X========X========X=== | 7 |
| chr03 | - | 36036711 | ATTTCAAATTTCAGTTACTTCTAAATTGTATT | =X=XX==X==X========X========X=== | 7 |
| chr04 | - | 17893572 | ATCACAAGTTTTAGTTATTTATAAATTGTATT | =X==X=====XX=====X=X========X=== | 7 |
| chr04 | - | 20265933 | ATTTCAAATTTCAGTTACTTCTAAATTGTATT | =X=XX==X==X========X========X=== | 7 |
| chr04 | - | 20666471 | ATTTCAAATTTCAGTTACTTCTAAATTGTATT | =X=XX==X==X========X========X=== | 7 |
| chr04 | - | 20666975 | ATTTTAAATTTCAGTTACTTCTAAATTGTATT | =X=XX==X==X========X========X=== | 7 |
| chr05 | - | 1735957 | AAGGGAAGATCCATATACCATCAAATTGTAGT | ===X======X==X====X==X======X=X= | 7 |
| chr05 | - | 25838709 | ATTTCAAATTTCAGTTACTTCTAAATTGTATT | =X=XX==X==X========X========X=== | 7 |
| chr06 | - | 6930826 | AAAAAAAGCTGCAGCTATTAACAAATGAGAGC | ====X============X===X=====X==XX | 6 |
| chr06 | - | 7492325 | AAAAGTAGATGGAGGGAGTACTAATTAGAAAT | =====X=====X===X=X======X===X=X= | 7 |
| chr06 | - | 9307802 | ATTTCAAATTTCAGTTACTTCTAAATTGTATT | =X=XX==X==X========X========X=== | 7 |
| chr06 | - | 20594939 | ATTACGAATTTCAGTTACTTATAAATTGTATT | =X==XX=X==X========X========X=== | 7 |
| chr06 | - | 24783660 | AATAGTAGAATCAGCTTCTACTATATCGTATC | =====X===XX=====X======X====X==X | 7 |
| chr06 | - | 28241160 | ATTTTAAATTTCAGTTACTTCTAAATTGTATT | =X=XX==X==X========X========X=== | 7 |
| chr07 | - | 5662145 | TATAGATCGATCAGTTAATATTTAATGGGATT | X=====XX=XX======X====X========= | 7 |
| chr07 | - | 25383171 | AGCAGCAGCTGCAGATATTACGAAGTATGCTT | =X===X===========X===X==X==X=X== | 7 |
| chr07 | - | 29239352 | ATTTCAAATTTCAGTTACTTCTAAATTGTATT | =X=XX==X==X========X========X=== | 7 |
| chr08 | - | 4302371 | ATTTCAAATTTCAGTTACTTCTAAATTGTATT | =X=XX==X==X========X========X=== | 7 |
| chr08 | - | 15512649 | ATTTCAAATTTCAGTTACTTCTAAATTGTATT | =X=XX==X==X========X========X=== | 7 |
| chr08 | - | 15513243 | ATTTCAAATTTCAGTTACTTCTAAATTGTATT | =X=XX==X==X========X========X=== | 7 |
| chr08 | - | 25339348 | ATTACAAATTTCAGTTAATTTTAAATTGTATT | =X==X==X==X======X=X========X=== | 7 |
| chr08 | - | 27110316 | AAATTAAGATAAAGATACTAGTAAATATTAAT | ===XX=====XX===============XX=X= | 7 |
| chr08 | - | 27871080 | ATTTTAAATTTCAGTTACTTCTAAATTGTATT | =X=XX==X==X========X========X=== | 7 |
| chr09 | - | 7157847 | AGAAGAAGTTGTAGTTGCTTTTATGTGGGATC | =X=========X====X==X===XX======X | 7 |
| chr09 | - | 12152866 | ATTTCAAATTTCAGTTACTTCTAAATTGTATT | =X=XX==X==X========X========X=== | 7 |
| chr09 | - | 16126378 | ATTTCAAATTTCAGTTACTTCTAAATTGTATT | =X=XX==X==X========X========X=== | 7 |
| chr09 | - | 17398245 | ATCAAAAGTTTTAGTTATTTATAAATTGTATT | =X==X=====XX=====X=X========X=== | 7 |
| chr10 | - | 5758283 | ATTTCAAATTTCAGTTACTTCTAAATTGTATT | =X=XX==X==X========X========X=== | 7 |
| chr10 | - | 16915064 | ATATGGAATTTCAGTTACTTCTAAATTGTATT | =X=X=X=X==X========X========X=== | 7 |

(C) OsDL (continued)

| chromosome | strand | start | sequence | Mismatch sites (X) | No. of mismatch |
|---|---|---|---|---|---|
| chr11 | - | 6435886 | ATTTCAAATTTCAGTTACTTCTAAATTGTATT | =X=XX==X==X========X========X=== | 7 |
| chr11 | - | 21573290 | AATTCGAATTTCAGTTACTTCTAAATTGTATT | ===XXX=X==X========X========X=== | 7 |
| chr11 | - | 26316052 | ATTTCAAATTTCAGTTACTTCTAAATTGTATT | =X=XX==X==X========X========X=== | 7 |
| chr11 | - | 27346599 | ATTTCGAATTTCAGTTACTAATAAATTGTATT | =X=XXX=X==X=================X=== | 7 |
| chr12 | - | 3806167 | AAGAGGAGTGGAAGATGCTAGTTAATTGTATG | =====X===X=X====X=====X=====X==X | 7 |
| chr12 | - | 22311379 | ATTCCAAGTTTCAGCTAATTTTAAATTGTATT | =X=XX=====X======X=X========X=== | 7 |
| chr12 | - | 25209942 | ATTTCAAATTTCAGTTACTTCTAAATTGTATT | =X=XX==X==X========X========X=== | 7 |

(D) OsWx (7-nt mismatch with potential PAM)

| chromosome | strand | start | sequence | Mismatch sites (X) | No. of mismatch |
|---|---|---|---|---|---|
| chr10 | + | 18401035 | TAGGCAATCGTTCTCGCCGTGGTAGCCTCCTC | X=X===X=======XX==========X====X | 7 |
| chr01 | - | 22388955 | TACGACGCCATCATGGTAAAGCCCAAAGCTTG | ==========X==X=====X=X=X=====X=X | 7 |

**Supplementary Table S3**. Oligonucleotides used in this study.

| | | |
|---|---|---|
| For vector construction | | |
| attB1 f | GGGGACAAGTTTGTACAAAAAAGCAGGCTTAGGCGCGCCAAGCT | Construction of pDONR221(L1-L4)ZmUbiCas5Tpea3A and pDONR221(L1-L4)ZmUbiCas6Tpea3A |
| attB4 r | GGGGACAACTTTGTATAGAAAAGTTGGGTGTTAATTAAGGTACCAAAGCCTATACTGTACT | |
| attP4 f | GGGGACAACTTTTCTATACAAAGTTGTAGGCGCGCCAAGCT | Construction of pDONR221(R4-R3)ZmUbiCas3Tpea3A and pDONR221(R4-R3)ZmUbiCas8Tpea3A |
| attP3 r | GGGGACAACTTTATTATACAAAGTTGTTTAATTAAGGTACCAAAGCCTATACTGTACT | |
| attB3 f | GGGGACAACTTTGTATAATAAAGTTGTAGGCGCGCCAAGCT | Construction of DONR221(L3-L2)ZmUbiCas7Tpea3A and pDONR221(L3-L2)ZmUbiCas11Tpea3A |
| attB2 r | GGGGACCACTTTGTACAAGAAAGCTGGGTTTTAATTAAGGTACCAAAGCCTATACTGTACT | |
| P35S-omega f1 | ATTTGGAGAGGCCGGTCTAGAGTATTTTTAC | Construction of pDONR221(L1-L4) 2x35SΩint Cas758 T17.3 and pDONR221(R4-R3)ZmUbiCas1163Tpea3A |
| RcCAT r2 | CCGTCGGCGGTCCTCTTGGCTCTGTAACTATCATCATCATCATAGACACACGAAATAAAG | |
| bpNLS f1 | GCCAAGAGGACCGCCGACGG | |
| T2A-NLS r1 | TTTTCCTCCACGTCCCCGCATGTTAGAAGACTTCCCCTGCCCTCGCCGGAGCCCTCAACCTTTCTTTTCTTCTTTGGGCTC | |
| T2A-NLS f1 | TGCGGGGACGTGGAGGAAAATCCCGGCCCAATGGCCAAGAGGACCGCC | |
| T2A-NLS r2 | TGTTAGAAGACTTCCCCTGCCCTCGCCGGAGCCCTCAACCTTCCGTTTCTTCTTTGGGCTC | |
| T2A-NLS f2 | GCAGGGGAAGTCTTCTAACATGCGGGGACGTGGAGGAAAATCCCGGCCCAATGGCCAAGAGGACCGCC | |
| T17.3 r1 | TTTTAGATTAGCCCAGAGCTCTCACTCAACCTTCCGTTTCTTCTTTGG | |
| ADHUTR-NLS f1 | AAAAGAGGGGGATTAACTAGATGGCCAAGAGGACCGCC | |
| T2A-NLS r3 | TGTTAGAAGACTTCCCCTGCCCTCGCCGGAGCCTTCGACCTTCCTCTTTTTCTTTGGGCTC | |
| Tpea3A r5 | AGCTGGGAGGCCTGGATCGG | |
| crOsPDS | ACCGGTTATGCATGCCTATTGTTGACCATGTTCGCT | Cloning into pOsU6Cas3 gRNA |
| | ACACAGCGAACATGGTCAACAATAGGCATGCATAAC | |
| crOsDL | ACCGAATCCAATTTAGTAGTACCTGCATCTTCTGTT | |
| | ACACAACAGAAGATGCAGGTACTACTAAATTGGATT | |
| crOsWx | ACCGAATGCATTCGTTCTTACCGTGGTTGCGTCGTA | |
| | ACACTACGACGCAACCACGGTAAGAACGAATGCATT | |
| nptII f2 | ATCTATCTCTCTCGAGCTTTCGCAGATC | Construction of pZD202Km |
| nptII r2 | TATGGAGAAACTCGAGCTTGTCGATC | |
| gRNA PmeI f | AAACACTGATAGTTTCCAAGCTTATCGATACCGTCGAC | gRNA cloning into *Pme*I-digested binary vectors |
| gRNA PmeI r | TCCCGCCTTCAGTTTGGTACCGAGCTCGGATCCAC | |
| gRNA SpeI f | AATTAACACCACTAGAAGCTTATCGATACCGTCGAC | gRNA cloning into *Spe*I-digested binary vectors |
| gRNA SpeI r | TTGTGATATCACTAGGGTACCGAGCTCGGATCCAC | |

| | | |
|---|---|---|
| For molecular analysis | | |
| OsPDS | CTTCACGCAGACGCCAAGATCAAGC | PCR analysis |
| | TGCAGAGGTCGGCAAGGTTCACAG | |
| OsDL | GGCTAGACCTAGCGCCCAACCAACC | |
| | AGCAGATGGGAGGCGGTGTTTCTTC | |
| OsPDS (chr7 off-target) | CGGCGGAATGGATTGCATAGTGGTTGA | |
| | CCAGGCGTATAGCCGCGTACTGCACAA | |
| OsDL (chr1 off-target) | ATGTGGCAGCTCGCGCTTCTGGAAC | |
| | CTGGGCAGGCACTGCAGAGTCGTTG | |
| OsDL (chr6 off-target) | TCCACCCTGCAAATACCACCCGGGATA | |
| | CGCCACTCGTCGGCGGGCAGAAGTC | |
| OsWx | CCCACTGGCGAGCTACCTGAAGAAC | |
| | GACGACGGATGCAACAAGAGCGTGAGAC | |
| OsPDS | ACACTCCATCAGTAAGTGCAAAGTGTTTCACTGTTC | Sanger sequencing |
| OsWx | TCATGCGGCTCACCGGCATCAC | |
| OsPDS | CCCAATGGTGTGTGTGTCATCATCC | probe synthesis |
| | GCAACCAGTGAGTGCAAAGGGAG | |
| OsDL | CGATGGAGATGCAAGTGTCTGATGA | |
| | GCTCCATCTGCTTGAAGATGTCGAA | |

**Supplementary sequence.** Sequences of expression cassettes used in this study.

| | |
|---|---|
| Cas3 | TAGAGATAATGAGCATTGCATGTCTAAGTTATAAAAAATTACCACATATTTTTTTTGTCACACTTGTTTGAAGTGCAGTTTATCTATCTTTATACATATATTTAAACTTTACTCTACGAAT<br>AATATAATCTATAGTACTACAATAATATCAGTGTTTTAGAGAATCATATAAATGAACAGTTAGACATGGTCTAAAGGACAATTGAGTATTTTGACAACAGGACTCTACAGTTTTATCTTT<br>TTAGTGTGCATGTGTTCTCCTTTTTTTTTGCAAATAGCTTCACCTATATAATACTTCATCCATTTTATTAGTACATCCATTTAGGGTTTAGGGTTAATGGTTTTTATAGACTAATTTTTTTA<br>GTACATCTATTTTATTCTATTTTAGCCTCTAAATTAAGAAAACTAAAACTCTATTTTAGTTTTTTTATTTAATAATTTAGATATAAAATAGAATAAAATAAAGTGACTAAAAATTAAACAAAT<br>ACCCTTTAAGAAATTAAAAAAACTAAGGAAACATTTTTCTTGTTTCGAGTAGATAATGCCAGCCTGTTAAACGCCGTCGACGAGTCTAACGGACACCAACCAGCGAACCAGCAGCGT<br>CGCGTCGGGCCAAGCGAAGCAGACGGCACGGCATCTCTGTCGCTGCCTCTGGACCCCTCTCGAGAGTTCCGCTCCACCGTTGGACTTGCTCCGCTGTCGGCATCCAGAAATTGC<br>GTGGCGGAGCGGCAGACGTGAGCCGGCACGGCAGGCGGCCTCCTCCTCCTCTCACGGCACGGCAGCTACGGGGGATTCCTTTCCCACCGCTCCTTCGCTTTCCCTTCCTCGCC<br>CGCCGTAATAAATAGACACCCCCTCCACACCCTCTTTCCCCAACCTCGTGTTGTTCGGAGCGCACACACACACAACCAGATCTCCCCCAAATCCACCCGTCGGCACCTCCGCTTC<br>AAGGTACGCCGCTCGTCCTCCCCCCCCCCCCCTCTCTACCTTCTCTAGATCGGCGTTCCGGTCCATGGTTAGGGCCCGGTAGTTCTACTTCTGTTCATGTTTGTGTTAGATCCGTG<br>TTTGTGTTAGATCCGTGCTGCTAGCGTTCGTACACGGATGCGACCTGTACGTCAGACACGTTCTGATTGCTAACTTGCCAGTGTTTCTCTTTGGGGAATCCTGGGATGGCTCTAGC<br>CGTTCCGCAGACGGGATCGATTTCATGATTTTTTTTGTTTCGTTGCATAGGGTTTGGTTTGCCCTTTTCCTTTATTTCAATATATGCCGTGCACTTGTTTGTCGGGTCATCTTTTCATG<br>CTTTTTTTTGTCTTGGTTGTGATGATGTGGTCTGGTTGGGCGGTCGTTCTAGATCGGAGTAGAATTCTGTTTCAAACTACCTGGTGGATTTATTAATTTTGGATCTGTATGTGTGTGC<br>CATACATATTCATAGTTACGAATTGAAGATGATGGATGGAAATATCGATCTAGGATAGGTATACATGTTGATGCGGGTTTTACTGATGCATATACAGAGATGCTTTTTGTTCGCTTGG<br>TTGTGATGATGTGGTGTGGTTGGGCGGTCGTTCATTCGTTCTAGATCGGAGTAGAATACTGTTTCAAACTACCTGGTGTATTTATTAATTTTGGAACTGTATGTGTGTGTCATACATC<br>TTCATAGTTACGAGTTTAAGATGGATGGAAATATCGATCTAGGATAGGTATACATGTTGATGTGGGTTTTACTGATGCATATACATGATGGCATATGCAGCATCTATTCATATGCTCTA<br>ACCTTGAGTACCTATCTATTATAATAAACAAGTATGTTTTATAATTATTTTGATCTTGATATACTTGGATGATGGCATATGCAGCAGCTATATGTGGATTTTTTTAGCCCTGCCTTCATA<br>CGCTATTTATTTGCTTGGTACTGTTTCTTTTGTCGATGCTCACCCTGTTGTTTGGTGTTACTTCTGCAGGTCGACTCTAGAGAATTCCAAGCAACGAACTGCGAGTGATTCAAGAAAA<br>AAGAAAACCTGAGCTTTCGATCTCTACGGAGTGGTTTCTTGTTCTTTGAAAAAGAGGGGGATTAACTAGTATGGCCAAGAGGACCGCCGACGGCTCTGAGTTTGAGTCCCCGAAGA<br>AGAAGCGCAAGGTCGAGGAACCGTTCAAGTACATCTGCCACTACTGGGGCAAGTCCTCCAAGAGCCTCACCAAGGGCAACGATATCCACCTCCTCATCTACCACTGCCTCGATGT<br>TGCCGCCGTTGCCGATTGCTGGTGGGATCAATCTGTGGTGCTCCAGAACACCTTCTGCCGCAACGAGATGCTCTCCAAGCAACGCGTTAAGGCCTGGCTCCTCTTCTTCATTGCC<br>CTCCACGACATCGGCAAGTTCGACATCCGCTTCCAGTACAAGTCCGCCGAGTCCTGGCTCAAGCTCAACCCAGCTACACCAAGCCTCAACGGCCCGTCCACACAAATGTGCCGC<br>AAGTTCAATCATGGCGCCGCTGGCCTCTACTGGTTCAACCAGGATTCTCTCAGCGAGCAGTCCCTCGGCGACTTCTTCTCCTTCTTCGATGCCGCGCCACATCCGTACGAGAGCT<br>GGTTCCCATGGGTTGAAGCCGTGACAGGCCACCATGGCTTCATCCTCCATAGCCAGGACCAGGACAAGTCCAGGTGGGAGATGCCAGCTTCTCTCGCCTCTTACGCCGCGCAGG<br>ATAAGCAAGCTCGCGAGGAATGGATCTCCGTGCTCGAGGCTCTGTTCCTCACACCAGCCGGCCTCTCCATCAACGACATTCCGCCAGATTGCTCCTCTCTCCTCGCTGGCTTTTG<br>CTCCCTCGCTGATTGGCTTGGCTCCTGGACCACCACCAACACCTTTCTGTTCAACGAGGACGCCCCGTCCGACATCAATGCCCTCAGGACCTACTTCCAGGACAGGCAGCAAGAT<br>GCCTCCAGGGTGCTCGAACTCTCTGGCCTCGTTTCCAACAAGCGCTGCTACGAGGGTGTGCATGCGCTCCTCGATAACGGCTACCAGCCAAGACAGCTCCAGGTCCTCGTGGAT<br>GCCCTTCCAGTTGCTCCAGGCCTCACAGTGATTGAGGCTCCAACAGGCTCCGGCAAGACAGAGACAGCCCTCGCCTATGCCTGGAAGCTCATCGATCAGCAGATCGCCGACTCC<br>GTGATCTTCGCGCTCCCAACACAAGCCACCGCCAATGCCATGCTCACCAGGATGGAAGCCTCTGCCAGCCACCTCTTCAGCTCCCCGAATCTCATTCTCGCCCACGGCAACTCC<br>CGCTTCAACCACCTCTTTCAGTCCATCAAGTCCCGCGCCATCACCGAGCAAGGCCAAGAAGAGGCTTGGGTGCAGTGCTGCCAATGGCTCTCCCAGTCCAACAAGAAGGTTTTCC<br>TCGGCCAGATCGGCGTGTGCACCATTGACCAGGTGCTCATCTCTGTGCTCCCGGTGAAGCACAGATTCATTCGCGGCCTTGGCATCGGCCGCTCAGTGCTCATTGTGGATGAGG<br>TGCACGCCTACGATACCTACATGAACGGCCTTCTCGAGGCCGTGCTCAAGGCCCAAGCCGATGTTGGCGGCTCTGTGATCCTCCTCTCTGCGACACTCCCAATGAAGCAGAAGC<br>AAAAGCTCCTCGACACCTACGGCCTCCACACCGATCCAGTCGAGAACAACTCCGCCTATCCGCTCATCAATTGGAGGGGCGTTAACGGCGCCCAGAGGTTCGATCTCCTTGCGC<br>ATCCAGAGCAGCTCCCGCCAAGGTTCTCTATCCAGCCTGAGCCAATCTGCCTCGCCGACATGCTCCCAGATCTCACCATGCTCGAGCGCATGATCGCCGCTGCTAATGCCGGTG<br>CTCAAGTGTGCCTCATCTGCAACCTCGTCGATGTGGCCCAGGTGTGCTACCAGAGGCTCAAAGAGCTGAACAACACCCAGGTGGACATCGACCTCTTCCACGCCAGGTTCACCCT<br>CAACGACCGCCGCGAGAAAGAGAACCGCGTCATCTCCAACTTCGGCAAGAACGGCAAGAGGAACGTGGGCAGAATCCTCGTGGCCACACAGGTGGTGGAACAGTCTCTCGACG<br>TGGACTTCGACTGGCTCATCACCCAACACTGCCCAGCCGACCTGCTCTTTCAAAGACTCGGCAGGCTCCACCGCCACCACAGAAAGTACAGACCAGCGGGCTTCGAGATCCCGG<br>TCGCCACAATTTTGCTGCCGGATGGCGAAGGCTACGGCAGGCACGAGCATATCTACTCCAACGTGAGGGTCATGTGGCGCACCCAGCAGCACATCGAGGAACTCAATGGCGCCA<br>GCCTGTTCTTCCCGGATGCCTACAGACAGTGGCTCGACTCCATCTACGACGACGCCGAGATGGATGAGCCTGAGTGGGTTGGCAACGGCATGGACAAGTTCGAGTCTGCCGAGT<br>GCGAGAAGAGGTTCAAGGCTCGCAAGGTGCTGCAGTGGGCCGAAGAGTACTCCCTCCAGGACAACGACGAGACAATCCTCGCCGTGACACGCGACGGCGAAATGTCTCTCCCA<br>CTCCTTCCGTACGTGCAGACCTCTTCTGGCAAGCAACTCCTCGACGGCCAGGTGTACGAGGATCTCTCTCATGAGCAGCAGTACGAGGCCCTCGCGCTCAATAGAGTGAACGTG<br>CCGTTTACCTGGAAGCGCTCCTTCAGCGAGGTGGTCGATGAAGATGGCCTCCTCTGGCTCGAGGGCAAGCAGAATCTTGATGGCTGGGTGTGGCAGGGCAACTCCATCGTGATC<br>ACCTACACAGGCGACGAGGGCATGACCAGGGTGATCCCAGCCAATCCAAAGAAGAGAACGGCGGACGGCAGCGAGTTCGAGAGCCCAAAGAAAAAGCGGAAGGTTGAGTGAGA<br>GCTCCGATCCAGGCCTCCCAGCTTTCGTCCGTATCATCGGTTTCGACAACGTTCGTCAAGTTCAATGCATCAGTTTCATTGCCCACACACCAGAATCCTACTAAGTTTGAGTATTAT<br>GGCATTGGAAAAGCTGTTTTCTTCTATCATTTGTTCTGCTTGTAATTTACTGTGTTCTTTCAGTTTTTGTTTTCGGACATCAAAATGCAAATGGATGGATAAGAGTTAATAAATGATATG<br>GTCCTTTTGTTCATTCTCAAATTATTATTATCTGTTGTTTTTACTTTAATGGGTTGAATTTAAGTAAGAAAGGAACTAACAGTGTGATATTAAGGTGCAATGTTAGACATATAAAACAGT<br>CTTTCACCTCTCTTTGGTTATGTCTTGAATTGGTTTGTTTCTTCACTTATCTGTGTAATCAAGTTTACTATGAGTCTATGATCAAGTAATTATGCAATCAAGTTAAGTACAGTATAGGCT<br>TT (ZmUbi promoter, OsADH 5’UTR, Cas3 with bpNLS, pea3A terminator) |
| Cas5 | TAGAGATAATGAGCATTGCATGTCTAAGTTATAAAAAATTACCACATATTTTTTTTGTCACACTTGTTTGAAGTGCAGTTTATCTATCTTTATACATATATTTAAACTTTACTCTACGAAT<br>AATATAATCTATAGTACTACAATAATATCAGTGTTTTAGAGAATCATATAAATGAACAGTTAGACATGGTCTAAAGGACAATTGAGTATTTTGACAACAGGACTCTACAGTTTTATCTTT<br>TTAGTGTGCATGTGTTCTCCTTTTTTTTTGCAAATAGCTTCACCTATATAATACTTCATCCATTTTATTAGTACATCCATTTAGGGTTTAGGGTTAATGGTTTTTATAGACTAATTTTTTTA<br>GTACATCTATTTTATTCTATTTTAGCCTCTAAATTAAGAAAACTAAAACTCTATTTTAGTTTTTTTATTTAATAATTTAGATATAAAATAGAATAAAATAAAGTGACTAAAAATTAAACAAAT<br>ACCCTTTAAGAAATTAAAAAAACTAAGGAAACATTTTTCTTGTTTCGAGTAGATAATGCCAGCCTGTTAAACGCCGTCGACGAGTCTAACGGACACCAACCAGCGAACCAGCAGCGT<br>CGCGTCGGGCCAAGCGAAGCAGACGGCACGGCATCTCTGTCGCTGCCTCTGGACCCCTCTCGAGAGTTCCGCTCCACCGTTGGACTTGCTCCGCTGTCGGCATCCAGAAATTGC<br>GTGGCGGAGCGGCAGACGTGAGCCGGCACGGCAGGCGGCCTCCTCCTCCTCTCACGGCACGGCAGCTACGGGGGATTCCTTTCCCACCGCTCCTTCGCTTTCCCTTCCTCGCC<br>CGCCGTAATAAATAGACACCCCCTCCACACCCTCTTTCCCCAACCTCGTGTTGTTCGGAGCGCACACACACACAACCAGATCTCCCCCAAATCCACCCGTCGGCACCTCCGCTTC<br>AAGGTACGCCGCTCGTCCTCCCCCCCCCCCCCTCTCTACCTTCTCTAGATCGGCGTTCCGGTCCATGGTTAGGGCCCGGTAGTTCTACTTCTGTTCATGTTTGTGTTAGATCCGTG<br>TTTGTGTTAGATCCGTGCTGCTAGCGTTCGTACACGGATGCGACCTGTACGTCAGACACGTTCTGATTGCTAACTTGCCAGTGTTTCTCTTTGGGGAATCCTGGGATGGCTCTAGC<br>CGTTCCGCAGACGGGATCGATTTCATGATTTTTTTTGTTTCGTTGCATAGGGTTTGGTTTGCCCTTTTCCTTTATTTCAATATATGCCGTGCACTTGTTTGTCGGGTCATCTTTTCATG<br>CTTTTTTTTGTCTTGGTTGTGATGATGTGGTCTGGTTGGGCGGTCGTTCTAGATCGGAGTAGAATTCTGTTTCAAACTACCTGGTGGATTTATTAATTTTGGATCTGTATGTGTGTGC<br>CATACATATTCATAGTTACGAATTGAAGATGATGGATGGAAATATCGATCTAGGATAGGTATACATGTTGATGCGGGTTTTACTGATGCATATACAGAGATGCTTTTTGTTCGCTTGG<br>TTGTGATGATGTGGTGTGGTTGGGCGGTCGTTCATTCGTTCTAGATCGGAGTAGAATACTGTTTCAAACTACCTGGTGTATTTATTAATTTTGGAACTGTATGTGTGTGTCATACATC<br>TTCATAGTTACGAGTTTAAGATGGATGGAAATATCGATCTAGGATAGGTATACATGTTGATGTGGGTTTTACTGATGCATATACATGATGGCATATGCAGCATCTATTCATATGCTCTA<br>ACCTTGAGTACCTATCTATTATAATAAACAAGTATGTTTTATAATTATTTTGATCTTGATATACTTGGATGATGGCATATGCAGCAGCTATATGTGGATTTTTTTAGCCCTGCCTTCATA<br>CGCTATTTATTTGCTTGGTACTGTTTCTTTTGTCGATGCTCACCCTGTTGTTTGGTGTTACTTCTGCAGGTCGACTCTAGAGAATTCCAAGCAACGAACTGCGAGTGATTCAAGAAAA<br>AAGAAAACCTGAGCTTTCGATCTCTACGGAGTGGTTTCTTGTTCTTTGAAAAAGAGGGGGATTAACTAGTATGGCCAAGAGGACCGCCGACGGCTCTGAGTTTGAGTCCCCGAAGA<br>AGAAGCGCAAGGTCGAGCGCTCCTACCTCATCCTTAGACTCGCTGGCCCAATGCAGGCTTGGGGCCAGCCAACATTTGAAGGCACCAGACCAACCGGCAGGTTCCCAACAAGAT<br>CTGGCCTCCTTGGCCTTCTCGGCGCTTGCCTTGGCATCCAGAGGGATGATACCTCCAGCCTCCAAGCGCTCTCTGAGTCTGTGCAATTCGCCGTGAGGTGCGACGAGCTGATCC<br>TCGATGATAGACGCGTGTCCGTGACCGGCCTCAGGGATTACCATACAGTTCTTGGCGCTCGCGAGGACTACAGGGGCCTTAAGTCCCACGAGACAATCCAGACCTGGCGCGAGT<br>ACCTCTGCGACGCCTCTTTCACAGTTGCCCTCTGGCTCACCCCACACGCCACAATGGTGATCTCCGAGCTTGAGAAGGCCGTGCTCAAGCCGAGGTACACACCTTATCTTGGCAG<br>GCGCAGCTGCCCGCTCACACATCCACTGTTTCTCGGCACATGCCAGGCCTCCGATCCTCAAAAGGCCCTCCTCAACTACGAGCCGGTTGGCGGCGATATCTACTCCGAGGAATC<br>CGTGACAGGCCACCACCTCAAGTTCACCGCTAGGGACGAGCCGATGATCACACTCCCAAGACAATTCGCCAGCCGCGAGTGGTACGTGATCAAAGGCGGCATGGACGTGTCCCA<br>GAAGAGAACAGCTGACGGCAGCGAGTTCGAGAGCCCAAAGAAGAAACGGAAGGTTGAGTGAGAGCTCCGATCCAGGCCTCCCAGCTTTCGTCCGTATCATCGGTTTCGACAACG<br>TTCGTCAAGTTCAATGCATCAGTTTCATTGCCCACACACCAGAATCCTACTAAGTTTGAGTATTATGGCATTGGAAAAGCTGTTTTCTTCTATCATTTGTTCTGCTTGTAATTTACTGT<br>GTTCTTTCAGTTTTTGTTTTCGGACATCAAAATGCAAATGGATGGATAAGAGTTAATAAATGATATGGTCCTTTTGTTCATTCTCAAATTATTATTATCTGTTGTTTTTACTTTAATGGGT<br>TGAATTTAAGTAAGAAAGGAACTAACAGTGTGATATTAAGGTGCAATGTTAGACATATAAAACAGTCTTTCACCTCTCTTTGGTTATGTCTTGAATTGGTTTGTTTCTTCACTTATCTGT<br>GTAATCAAGTTTACTATGAGTCTATGATCAAGTAATTATGCAATCAAGTTAAGTACAGTATAGGCTTT (ZmUbi promoter, OsADH 5’UTR, Cas5 with bpNLS, pea3A terminator) |

| | |
|---|---|
| Cas6 | TAGAGATAATGAGCATTGCATGTCTAAGTTATAAAAAATTACCACATATTTTTTTTGTCACACTTGTTTGAAGTGCAGTTTATCTATCTTTATACATATATTTAAACTTTACTCTACGAATAA<br>TATAATCTATAGTACTACAATAATATCAGTGTTTTAGAGAATCATATAAATGAACAGTTAGACATGGTCTAAAGGACAATTGAGTATTTTGACAACAGGACTCTACAGTTTTATCTTTTTA<br>GTGTGCATGTGTTCTCCTTTTTTTTTGCAAATAGCTTCACCTATATAATACTTCATCCATTTTATTAGTACATCCATTTAGGGTTTAGGGTTAATGGTTTTTATAGACTAATTTTTTTAGTAC<br>ATCTATTTTATTCTATTTTAGCCTCTAAATTAAGAAAACTAAAACTCTATTTTAGTTTTTTTATTTAATAATTTAGATATAAAATAGAATAAAATAAAGTGACTAAAAATTAAACAAATACCCT<br>TTAAGAAATTAAAAAAACTAAGGAAACATTTTTCTTGTTTCGAGTAGATAATGCCAGCCTGTTAAACGCCGTCGACGAGTCTAACGGACACCAACCAGCGAACCAGCAGCGTCGCGTC<br>GGGCCAAGCGAAGCAGACGGCACGGCATCTCTGTCGCTGCCTCTGGACCCCTCTCGAGAGTTCCGCTCCACCGTTGGACTTGCTCCGCTGTCGGCATCCAGAAATTGCGTGGCGG<br>AGCGGCAGACGTGAGCCGGCACGGCAGGCGGCCTCCTCCTCCTCTCACGGCACGGCAGCTACGGGGGATTCCTTTCCCACCGCTCCTTCGCTTTCCCTTCCTCGCCCGCCGTAAT<br>AAATAGACACCCCCTCCACACCCTCTTTCCCCAACCTCGTGTTGTTCGGAGCGCACACACACACAACCAGATCTCCCCCAAATCCACCCGTCGGCACCTCCGCTTCAAGGTACGCC<br>GCTCGTCCTCCCCCCCCCCCCCTCTCTACCTTCTCTAGATCGGCGTTCCGGTCCATGGTTAGGGCCCGGTAGTTCTACTTCTGTTCATGTTTGTGTTAGATCCGTGTTTGTGTTAGAT<br>CCGTGCTGCTAGCGTTCGTACACGGATGCGACCTGTACGTCAGACACGTTCTGATTGCTAACTTGCCAGTGTTTCTCTTTGGGGAATCCTGGGATGGCTCTAGCCGTTCCGCAGACG<br>GGATCGATTTCATGATTTTTTTTGTTTCGTTGCATAGGGTTTGGTTTGCCCTTTTCCTTTATTTCAATATATGCCGTGCACTTGTTTGTCGGGTCATCTTTTCATGCTTTTTTTTGTCTTGG<br>TTGTGATGATGTGGTCTGGTTGGGCGGTCGTTCTAGATCGGAGTAGAATTCTGTTTCAAACTACCTGGTGGATTTATTAATTTTGGATCTGTATGTGTGTGCCATACATATTCATAGTTA<br>CGAATTGAAGATGATGGATGGAAATATCGATCTAGGATAGGTATACATGTTGATGCGGGTTTTACTGATGCATATACAGAGATGCTTTTTGTTCGCTTGGTTGTGATGATGTGGTGTGG<br>TTGGGCGGTCGTTCATTCGTTCTAGATCGGAGTAGAATACTGTTTCAAACTACCTGGTGTATTTATTAATTTTGGAACTGTATGTGTGTGTCATACATCTTCATAGTTACGAGTTTAAGA<br>TGGATGGAAATATCGATCTAGGATAGGTATACATGTTGATGTGGGTTTTACTGATGCATATACATGATGGCATATGCAGCATCTATTCATATGCTCTAACCTTGAGTACCTATCTATTAT<br>AATAAACAAGTATGTTTTATAATTATTTTGATCTTGATATACTTGGATGATGGCATATGCAGCAGCTATATGTGGATTTTTTTAGCCCTGCCTTCATACGCTATTTATTTGCTTGGTACTG<br>TTTCTTTTGTCGATGCTCACCCTGTTGTTTGGTGTTACTTCTGCAGGTCGACTCTAGAGAATTCCAAGCAACGAACTGCGAGTGATTCAAGAAAAAAGAAAACCTGAGCTTTCGATCTC<br>TACGGAGTGGTTTCTTGTTCTTTGAAAAAGAGGGGGATTAACTAGTATGGCCAAGAGGACCGCCGACGGCTCTGAGTTTGAGTCCCCGAAGAAGAAGCGCAAGGTCGAGTACCTCT<br>CCAAGGTGATCATTGCCCGCGCTTGGTCCAGGGACCTCTACCAACTTCATCAAGGCCTCTGGCATCTCTTCCCGAATCGCCCTGATGCCGCCAGGGATTTCCTGTTCCATGTCGAG<br>AAGAGGAACACCCCAGAGGGCTGCCATGTGCTCCTCCAATCTGCCCAAATGCCAGTGTCTACCGCCGTGGCCACCGTGATCAAGACCAAGCAGGTTGAGTTCCAGCTCCAGGTGG<br>GCGTGCCACTCTATTTCAGGCTCAGGGCGAACCCGATCAAGACGATCCTCGACAACCAGAAGCGCCTCGACTCCAAGGGCAACATCAAGAGGTGCAGGGTGCCGCTCATCAAAGA<br>GGCCGAGCAAATTGCCTGGCTCCAGCGCAAGCTTGGCAATGCTGCCAGAGTCGAGGATGTGCACCCAATCTCTGAGCGCCCGCAATACTTCTCTGGCGACGGCAAGTCTGGCAAG<br>ATCCAGACCGTTTGCTTCGAGGGCGTGCTCACCATCAATGATGCCCCAGCGCTCATCGACCTCGTGCAGCAAGGCATTGGACCGGCTAAGTCTATGGGCTGCGGCCTCCTCTCACT<br>CGCTCCACTTAAGAGAACAGCGGACGGCAGCGAGTTCGAGAGCCCAAAGAAGAAAAGAAAGGTTGAGTGAGAGCTCCGATCCAGGCCTCCCAGCTTTCGTCCGTATCATCGGTTTC<br>GACAACGTTCGTCAAGTTCAATGCATCAGTTTCATTGCCCACACACCAGAATCCTACTAAGTTTGAGTATTATGGCATTGGAAAAGCTGTTTTCTTCTATCATTTGTTCTGCTTGTAATT<br>TACTGTGTTCTTTCAGTTTTTGTTTTCGGACATCAAAATGCAAATGGATGGATAAGAGTTAATAAATGATATGGTCCTTTTGTTCATTCTCAAATTATTATTATCTGTTGTTTTTACTTTAAT<br>GGGTTGAATTTAAGTAAGAAAGGAACTAACAGTGTGATATTAAGGTGCAATGTTAGACATATAAAACAGTCTTTCACCTCTCTTTGGTTATGTCTTGAATTGGTTTGTTTCTTCACTTATC<br>TGTGTAATCAAGTTTACTATGAGTCTATGATCAAGTAATTATGCAATCAAGTTAAGTACAGTATAGGCTTT (ZmUbi promoter, OsADH 5'UTR, Cas6 with bpNLS, pea3A terminator) |
| Cas7 | TAGAGATAATGAGCATTGCATGTCTAAGTTATAAAAAATTACCACATATTTTTTTTGTCACACTTGTTTGAAGTGCAGTTTATCTATCTTTATACATATATTTAAACTTTACTCTACGAATAA<br>TATAATCTATAGTACTACAATAATATCAGTGTTTTAGAGAATCATATAAATGAACAGTTAGACATGGTCTAAAGGACAATTGAGTATTTTGACAACAGGACTCTACAGTTTTATCTTTTTA<br>GTGTGCATGTGTTCTCCTTTTTTTTTGCAAATAGCTTCACCTATATAATACTTCATCCATTTTATTAGTACATCCATTTAGGGTTTAGGGTTAATGGTTTTTATAGACTAATTTTTTTAGTAC<br>ATCTATTTTATTCTATTTTAGCCTCTAAATTAAGAAAACTAAAACTCTATTTTAGTTTTTTTATTTAATAATTTAGATATAAAATAGAATAAAATAAAGTGACTAAAAATTAAACAAATACCCT<br>TTAAGAAATTAAAAAAACTAAGGAAACATTTTTCTTGTTTCGAGTAGATAATGCCAGCCTGTTAAACGCCGTCGACGAGTCTAACGGACACCAACCAGCGAACCAGCAGCGTCGCGTC<br>GGGCCAAGCGAAGCAGACGGCACGGCATCTCTGTCGCTGCCTCTGGACCCCTCTCGAGAGTTCCGCTCCACCGTTGGACTTGCTCCGCTGTCGGCATCCAGAAATTGCGTGGCGG<br>AGCGGCAGACGTGAGCCGGCACGGCAGGCGGCCTCCTCCTCCTCTCACGGCACGGCAGCTACGGGGGATTCCTTTCCCACCGCTCCTTCGCTTTCCCTTCCTCGCCCGCCGTAAT<br>AAATAGACACCCCCTCCACACCCTCTTTCCCCAACCTCGTGTTGTTCGGAGCGCACACACACACAACCAGATCTCCCCCAAATCCACCCGTCGGCACCTCCGCTTCAAGGTACGCC<br>GCTCGTCCTCCCCCCCCCCCCCTCTCTACCTTCTCTAGATCGGCGTTCCGGTCCATGGTTAGGGCCCGGTAGTTCTACTTCTGTTCATGTTTGTGTTAGATCCGTGTTTGTGTTAGAT<br>CCGTGCTGCTAGCGTTCGTACACGGATGCGACCTGTACGTCAGACACGTTCTGATTGCTAACTTGCCAGTGTTTCTCTTTGGGGAATCCTGGGATGGCTCTAGCCGTTCCGCAGACG<br>GGATCGATTTCATGATTTTTTTTGTTTCGTTGCATAGGGTTTGGTTTGCCCTTTTCCTTTATTTCAATATATGCCGTGCACTTGTTTGTCGGGTCATCTTTTCATGCTTTTTTTTGTCTTGG<br>TTGTGATGATGTGGTCTGGTTGGGCGGTCGTTCTAGATCGGAGTAGAATTCTGTTTCAAACTACCTGGTGGATTTATTAATTTTGGATCTGTATGTGTGTGCCATACATATTCATAGTTA<br>CGAATTGAAGATGATGGATGGAAATATCGATCTAGGATAGGTATACATGTTGATGCGGGTTTTACTGATGCATATACAGAGATGCTTTTTGTTCGCTTGGTTGTGATGATGTGGTGTGG<br>TTGGGCGGTCGTTCATTCGTTCTAGATCGGAGTAGAATACTGTTTCAAACTACCTGGTGTATTTATTAATTTTGGAACTGTATGTGTGTGTCATACATCTTCATAGTTACGAGTTTAAGA<br>TGGATGGAAATATCGATCTAGGATAGGTATACATGTTGATGTGGGTTTTACTGATGCATATACATGATGGCATATGCAGCATCTATTCATATGCTCTAACCTTGAGTACCTATCTATTAT<br>AATAAACAAGTATGTTTTATAATTATTTTGATCTTGATATACTTGGATGATGGCATATGCAGCAGCTATATGTGGATTTTTTTAGCCCTGCCTTCATACGCTATTTATTTGCTTGGTACTG<br>TTTCTTTTGTCGATGCTCACCCTGTTGTTTGGTGTTACTTCTGCAGGTCGACTCTAGAGAATTCCAAGCAACGAACTGCGAGTGATTCAAGAAAAAAGAAAACCTGAGCTTTCGATCTC<br>TACGGAGTGGTTTCTTGTTCTTTGAAAAAGAGGGGGATTAACTAGTATGGCCAAGAGGACCGCCGACGGCTCTGAGTTTGAGTCCCCGAAGAAGAAGCGCAAGGTCGAGTCCAACT<br>TCATCAACATCCACGTGCTCATCTCCCACTCGCCGTCCTGCCTCAACAGGGACGACATGAACATGCAGAAGGACGCCATCTTCGGCGGCAAGAGGCGCGTTAGAATTTCCAGCCAG<br>AGCCTCAAGCGCGCCATGAGGAAGTCTGGCTACTACGCCCAGAACATCGGCGAGTCTAGCCTCCGCACAATCCATCTCGCCCAGCTCAGAGATGTGCTCAGGCAAAAGCTCGGCG<br>AGCGCTTCGACCAGAAGATCATCGATAAGACCCTCGCGCTCCTCAGCGGCAAGTCTGTTGATGAGGCCGAGAAGATCTCCGCCGATGCCGTGACACCATGGGTGGTGGGCGAAAT<br>TGCCTGGTTCTGCGAGCAAGTTGCCAAGGCCGAGGCCGACAACCTCGACGACAAGAAGCTCCTCAAGGTCCTCAAAGAGGATATCGCCGCCATCCGCGTGAACCTCCAGCAAGGT<br>GTTGATATCGCTCTCTCCGGCAGGATGGCCACCTCTGGCATGATGACAGAGCTTGGCAAAGTGGACGGCGCCATGTCTATCGCCCATGCCATCACCACACACCAGGTGGACTCCG<br>ACATCGACTGGTTCACCGCCGTGGACGATCTCCAAGAACAAGGCTCTGCCCACCTCGGGACCCAAGAATTTTCTTCCGGCGTGTTCTACCGCTACGCCAACATCAACCTCGCGCAG<br>CTCCAAGAGAATCTCGGCGGAGCTTCTAGAGAGCAGGCCCTCGAGATTGCCACACACGTGGTGCATATGCTCGCCACAGAAGTGCCAGGCGCCAAGCAAAGAACCTACGCCGCCT<br>TCAATCCGGCCGACATGGTGATGGTGAACTTCTCCGATATGCCGCTCTCCATGGCCAACGCCTTCGAGAAGGCCGTGAAGGCCAAGGATGGCTTCCTCCAGCCATCCATCCAGGC<br>CTTCAACCAGTATTGGGACCGCGTGGCCAATGGCTACGGCCTTAATGGTGCCGCCGCTCAGTTCTCCCTCTCCGATGTGGATCCAATCACGGCCCAGGTGAAGCAGATGCCGACA<br>CTCGAGCAACTCAAGTCCTGGGTGCGCAACAATGGCGAGGCTAAGAGAACAGCCGATGGCAGCGAGTTCGAGAGCCCAAAGAAGAAAAGAAAGGTTGAGTGAGAGCTCCGATCCA<br>GGCCTCCCAGCTTTCGTCCGTATCATCGGTTTCGACAACGTTCGTCAAGTTCAATGCATCAGTTTCATTGCCCACACACCAGAATCCTACTAAGTTTGAGTATTATGGCATTGGAAAA<br>GCTGTTTTCTTCTATCATTTGTTCTGCTTGTAATTTACTGTGTTCTTTCAGTTTTTGTTTTCGGACATCAAAATGCAAATGGATGGATAAGAGTTAATAAATGATATGGTCCTTTTGTTCAT<br>TCTCAAATTATTATTATCTGTTGTTTTTACTTTAATGGGTTGAATTTAAGTAAGAAAGGAACTAACAGTGTGATATTAAGGTGCAATGTTAGACATATAAAACAGTCTTTCACCTCTCTTTG<br>GTTATGTCTTGAATTGGTTTGTTTCTTCACTTATCTGTGTAATCAAGTTTACTATGAGTCTATGATCAAGTAATTATGCAATCAAGTTAAGTACAGTATAGGCTTT (ZmUbi promoter,<br>OsADH 5'UTR, Cas7 with bpNLS, pea3A terminator) |

| | |
|---|---|
| Cas8 | TAGAGATAATGAGCATTGCATGTCTAAGTTATAAAAAATTACCACATATTTTTTTTGTCACACTTGTTTGAAGTGCAGTTTATCTATCTTTATACATATATTTAAACTTTACTCTACGAA<br>TAATATAATCTATAGTACTACAATAATATCAGTGTTTTAGAGAATCATATAAATGAACAGTTAGACATGGTCTAAAGGACAATTGAGTATTTTGACAACAGGACTCTACAGTTTTATCT<br>TTTTAGTGTGCATGTGTTCTCCTTTTTTTTTGCAAATAGCTTCACCTATATAATACTTCATCCATTTTATTAGTACATCCATTTAGGGTTTAGGGTTAATGGTTTTTATAGACTAATTTT<br>TTTAGTACATCTATTTTATTCTATTTTAGCCTCTAAATTAAGAAAACTAAAACTCTATTTTAGTTTTTTTATTTAATAATTTAGATATAAAATAGAATAAAATAAAGTGACTAAAAATTAAA<br>CAAATACCCTTTAAGAAATTAAAAAAACTAAGGAAACATTTTTCTTGTTTCGAGTAGATAATGCCAGCCTGTTAAACGCCGTCGACGAGTCTAACGGACACCAACCAGCGAACCAG<br>CAGCGTCGCGTCGGGCCAAGCGAAGCAGACGGCACGGCATCTCTGTCGCTGCCTCTGGACCCCTCTCGAGAGTTCCGCTCCACCGTTGGACTTGCTCCGCTGTCGGCATCCA<br>GAAATTGCGTGGCGGAGCGGCAGACGTGAGCCGGCACGGCAGGCGGCCTCCTCCTCCTCTCACGGCACGGCAGCTACGGGGGATTCCTTTCCCACCGCTCCTTCGCTTTCCC<br>TTCCTCGCCCGCCGTAATAAATAGACACCCCCTCCACACCCTCTTTCCCCAACCTCGTGTTGTTCGGAGCGCACACACACACACAACCAGATCTCCCCCAAATCCACCCGTCGGCA<br>CCTCCGCTTCAAGGTACGCCGCTCGTCCTCCCCCCCCCCCCCTCTCTACCTTCTCTAGATCGGCGTTCCGGTCCATGGTTAGGGCCCGGTAGTTCTACTTCTGTTCATGTTTGT<br>GTTAGATCCGTGTTTGTGTTAGATCCGTGCTGCTAGCGTTCGTACACGGATGCGACCTGTACGTCAGACACGTTCTGATTGCTAACTTGCCAGTGTTTCTCTTTGGGGAATCCTG<br>GGATGGCTCTAGCCGTTCCGCAGACGGGATCGATTTCATGATTTTTTTTGTTTCGTTGCATAGGGTTTGGTTTGCCCTTTTCCTTTATTTCAATATATGCCGTGCACTTGTTTGTCG<br>GGTCATCTTTTCATGCTTTTTTTTGTCTTGGTTGTGATGATGTGGTCTGGTTGGGCGGTCGTTCTAGATCGGAGTAGAATTCTGTTTCAAACTACCTGGTGGATTTATTAATTTTGGA<br>TCTGTATGTGTGTGCCATACATATTCATAGTTACGAATTGAAGATGATGGATGGAAATATCGATCTAGGATAGGTATACATGTTGATGCGGGTTTTACTGATGCATATACAGAGATG<br>CTTTTTGTTCGCTTGGTTGTGATGATGTGGTGTGGTTGGGCGGTCGTTCATTCGTTCTAGATCGGAGTAGAATACTGTTTCAAACTACCTGGTGTATTTATTAATTTTGGAACTGTA<br>TGTGTGTGTCATACATCTTCATAGTTACGAGTTTAAGATGGATGGAAATATCGATCTAGGATAGGTATACATGTTGATGTGGGTTTTACTGATGCATATACATGATGGCATATGCAG<br>CATCTATTCATATGCTCTAACCTTGAGTACCTATCTATTATAATAAACAAGTATGTTTTATAATTATTTTGATCTTGATATACTTGGATGATGGCATATGCAGCAGCTATATGTGGATT<br>TTTTTAGCCCTGCCTTCATACGCTATTTATTTGCTTGGTACTGTTTCTTTTGTCGATGCTCACCCTGTTGTTTGGTGTTACTTCTGCAGGTCGACTCTAGAGAATTCCAAGCAACGA<br>ACTGCGAGTGATTCAAGAAAAAAGAAAACCTGAGCTTTCGATCTCTACGGAGTGGTTTCTTGTTCTTTGAAAAAGAGGGGGATTAACTAGTATGGCCAAGAGGACCGCCGACGGC<br>TCTGAGTTTGAGTCCCCGAAGAAGAAGCGCAAGGTCGAGAACCTCCTCATCGACAACTGGATCCCAGTGAGGCCTAGGAACGGCGGCAAGGTGCAGATCATCAACCTCCAGAG<br>CCTCTACTGCTCCAGGGATCAATGGCGCCTCTCACTCCCGAGGGACGATGGAACTCGCTGCTCTCGCTCTCCTCGTGTGCATCGGCCAGATTATTGCCCCAGCCAAGGACG<br>ACGTCGAGTTCCGCCATAGGATCATGAACCCGCTCACCGAGGACGAGTTCCAGCAGCTCATTGCCCCGTGGATCGACATGTTCTACCTCAACCACGCGGAGCACCCGTTCATG<br>CAGACAAAGGGCGTGAAGGCCAACGACGTGACCCCGATGGAAAAGCTCCTTGCTGGCGTTTCCGGCGCCACCAATTGCGCCTTCGTTAACCAGCCAGGCCAAGGCGAAGCTC<br>TTTGCGGAGGCTGTACTGCGATCGCCCTCTTCAACCAGGCCAATCAAGCTCCAGGCTTCGGCGGAGGCTTTAAGTCTGGCCTTAGAGGCGGCACCCCAGTGACCACATTCGTG<br>AGGGGCATTGACCTCAGGTCCACCGTGCTCCTCAACGTGCTCACACTCCCAAGGCTCCAGAAGCAGTTCCCGAACGAGTCCCACACCGAGAACCAGCCGACCTGGATCAAGCC<br>GATCAAGTCCAACGAGAGCATCCCGGCCTCTTCCATCGGCTTCGTTAGGGGCCTCTTCTGGCAGCCAGCGCATATCGAGCTGTGCGATCCAATCGGAATCGGCAAGTGCTCCT<br>GCTGCGGCCAAGAGTCCAACCTCAGGTATACCGGCTTCCTGAAAGAGAAGTTCACCTTCACCGTGAACGGCCTCTGGCCGCATCCACATTCTCCATGCCTCGTGACCGTGAAG<br>AAGGGCGAAGTCGAGGAAAAGTTCCTCGCCTTCACCACAAGCGCCCCAAGCTGGACACAAATCTCCCGCGTGGTGGTGGACAAGATCATCCAGAACGAGAACGGCAACAGGGT<br>CGCCGCCGTGGTGAATCAGTTCCGCAATATCGCCCCACAGAGCCCGCTCGAGCTTATCATGGGCGGCTACCGCAACAACCAGGCGAGCATTCTTGAGCGCAGGCACGACGTG<br>CTCATGTTCAATCAAGGCTGGCAGCAGTACGGCAACGTGATCAACGAGATCGTCACCGTCGGCCTCGGCTACAAGACAGCCCTTAGAAAGGCCCTCTACACCTTCGCCGAGGG<br>CTTCAAGAACAAGGACTTCAAAGGCGCTGGCGTGAGCGTGCACGAGACAGCTGAGGCATTTCTACCGCCAGTCCGAGCTGCTCATCCCAGATGTGCTCGCCAACGTGAACT<br>TCAGCCAGGCGGATGAGGTGATCGCCGACCTCAGGGATAAGCTCCATCAGCTCTGCGAGATGCTGTTCAACCAGAGCGTGGCCCCATACGCGCACCATCCAAAGCTCATCTCT<br>ACCCTCGCTCTCGCCAGGGCGACACTCTACAAGCACCTGAGGGAACTCAAGCCGCAAGGCGGCCCATCTAACGGCAAGAGAACTGCTGATGGCTCCGAGTTCGAGAGCCCAA<br>AGAAGAAACGGAAGGTTGAGTGAGAGCTCCGATCCAGGCCTCCCAGCTTTCGTCCGTATCATCGGTTTCGACAACGTTCGTCAAGTTCAATGCATCAGTTTCATTGCCCACACAC<br>CAGAATCCTACTAAGTTTGAGTATTATGGCATTGGAAAAGCTGTTTTCTTCTATCATTTGTTCTGCTTGTAATTTACTGTGTTCTTTCAGTTTTTGTTTTCGGACATCAAAATGCAAAT<br>GGATGGATAAGAGTTAATAAATGATATGGTCCTTTTGTTCATTCTCAAATTATTATTATCTGTTGTTTTTACTTTAATGGGTTGAATTTAAGTAAGAAAGGAACTAACAGTGTGATATT<br>AAGGTGCAATGTTAGACATATAAAACAGTCTTTCACCTCTCTTTGGTTATGTCTTGAATTGGTTTGTTTCTTCACTTATCTGTGTAATCAAGTTTACTATGAGTCTATGATCAAGTAAT<br>TATGCAATCAAGTTAAGTACAGTATAGGCTTT (ZmUbi promoter, OsADH 5’UTR, Cas8 with bpNLS, pea3A terminator) |
| Cas11 | TAGAGATAATGAGCATTGCATGTCTAAGTTATAAAAAATTACCACATATTTTTTTTGTCACACTTGTTTGAAGTGCAGTTTATCTATCTTTATACATATATTTAAACTTTACTCTACGAA<br>TAATATAATCTATAGTACTACAATAATATCAGTGTTTTAGAGAATCATATAAATGAACAGTTAGACATGGTCTAAAGGACAATTGAGTATTTTGACAACAGGACTCTACAGTTTTATCT<br>TTTTAGTGTGCATGTGTTCTCCTTTTTTTTTGCAAATAGCTTCACCTATATAATACTTCATCCATTTTATTAGTACATCCATTTAGGGTTTAGGGTTAATGGTTTTTATAGACTAATTTT<br>TTTAGTACATCTATTTTATTCTATTTTAGCCTCTAAATTAAGAAAACTAAAACTCTATTTTAGTTTTTTTATTTAATAATTTAGATATAAAATAGAATAAAATAAAGTGACTAAAAATTAAA<br>CAAATACCCTTTAAGAAATTAAAAAAACTAAGGAAACATTTTTCTTGTTTCGAGTAGATAATGCCAGCCTGTTAAACGCCGTCGACGAGTCTAACGGACACCAACCAGCGAACCAG<br>CAGCGTCGCGTCGGGCCAAGCGAAGCAGACGGCACGGCATCTCTGTCGCTGCCTCTGGACCCCTCTCGAGAGTTCCGCTCCACCGTTGGACTTGCTCCGCTGTCGGCATCCA<br>GAAATTGCGTGGCGGAGCGGCAGACGTGAGCCGGCACGGCAGGCGGCCTCCTCCTCCTCTCACGGCACGGCAGCTACGGGGGATTCCTTTCCCACCGCTCCTTCGCTTTCCC<br>TTCCTCGCCCGCCGTAATAAATAGACACCCCCTCCACACCCTCTTTCCCCAACCTCGTGTTGTTCGGAGCGCACACACACACACAACCAGATCTCCCCCAAATCCACCCGTCGGCA<br>CCTCCGCTTCAAGGTACGCCGCTCGTCCTCCCCCCCCCCCCCTCTCTACCTTCTCTAGATCGGCGTTCCGGTCCATGGTTAGGGCCCGGTAGTTCTACTTCTGTTCATGTTTGT<br>GTTAGATCCGTGTTTGTGTTAGATCCGTGCTGCTAGCGTTCGTACACGGATGCGACCTGTACGTCAGACACGTTCTGATTGCTAACTTGCCAGTGTTTCTCTTTGGGGAATCCTG<br>GGATGGCTCTAGCCGTTCCGCAGACGGGATCGATTTCATGATTTTTTTTGTTTCGTTGCATAGGGTTTGGTTTGCCCTTTTCCTTTATTTCAATATATGCCGTGCACTTGTTTGTCG<br>GGTCATCTTTTCATGCTTTTTTTTGTCTTGGTTGTGATGATGTGGTCTGGTTGGGCGGTCGTTCTAGATCGGAGTAGAATTCTGTTTCAAACTACCTGGTGGATTTATTAATTTTGGA<br>TCTGTATGTGTGTGCCATACATATTCATAGTTACGAATTGAAGATGATGGATGGAAATATCGATCTAGGATAGGTATACATGTTGATGCGGGTTTTACTGATGCATATACAGAGATG<br>CTTTTTGTTCGCTTGGTTGTGATGATGTGGTGTGGTTGGGCGGTCGTTCATTCGTTCTAGATCGGAGTAGAATACTGTTTCAAACTACCTGGTGTATTTATTAATTTTGGAACTGTA<br>TGTGTGTGTCATACATCTTCATAGTTACGAGTTTAAGATGGATGGAAATATCGATCTAGGATAGGTATACATGTTGATGTGGGTTTTACTGATGCATATACATGATGGCATATGCAG<br>CATCTATTCATATGCTCTAACCTTGAGTACCTATCTATTATAATAAACAAGTATGTTTTATAATTATTTTGATCTTGATATACTTGGATGATGGCATATGCAGCAGCTATATGTGGATT<br>TTTTTAGCCCTGCCTTCATACGCTATTTATTTGCTTGGTACTGTTTCTTTTGTCGATGCTCACCCTGTTGTTTGGTGTTACTTCTGCAGGTCGACTCTAGAGAATTCCAAGCAACGA<br>ACTGCGAGTGATTCAAGAAAAAAGAAAACCTGAGCTTTCGATCTCTACGGAGTGGTTTCTTGTTCTTTGAAAAAGAGGGGGATTAACTAGTATGGCCAAGAGGACCGCCGACGGC<br>TCTGAGTTTGAGTCCCCGAAGAAGAAGCGCAAGGTCGAGGCCGACGAGATCGATGCCATGGCTCTTTACAGAGCGTGGCAGCAGCTCGACAATGGCTCTTGCGCTCAGATTCG<br>CAGGGTGTCCGAGCCAGATGAGCTGAGGGATATTCCGGCCTTCTACAGGCTCGTGCAGCCATTCGGCTGGGAGAACCCAAGACATCAGCAGGCGCTCCTCCGCATGGTTTTCT<br>GCCTTTCCGCCGGCAAGAACGTGATCCGCCACCAGGATAAGAAGTCCGAGCAGACCACCGGCATTTCTCTCGGCAGAGCCCTTGCTAACTCCGGCAGGATCAATGAGCGCCG<br>CATCTTCCAGCTCATCAGGGCCGATAGGACAGCCGACATGGTTCAGCTCAGACGCCTCCTCACACATGCCGAGCCAGTTCTCGATTGGCCACTCATGGCCAGGATGCTCACAT<br>GGTGGGGAAAGAGAGAGCGCCAGCAGCTCCTTGAGGACTTCGTGCTCACCACCAACAAGAACGCCAAGCGCACAGCTGACGGCAGCGAGTTTGAGAGCCCAAAGAAAAAGAG<br>GAAGGTCGAAGGCTCCGGCTGAGAGCTCCGATCCAGGCCTCCCAGCTTTCGTCCGTATCATCGGTTTCGACAACGTTCGTCAAGTTCAATGCATCAGTTTCATTGCCCACACAC<br>CAGAATCCTACTAAGTTTGAGTATTATGGCATTGGAAAAGCTGTTTTCTTCTATCATTTGTTCTGCTTGTAATTTACTGTGTTCTTTCAGTTTTTGTTTTCGGACATCAAAATGCAAAT<br>GGATGGATAAGAGTTAATAAATGATATGGTCCTTTTGTTCATTCTCAAATTATTATTATCTGTTGTTTTTACTTTAATGGGTTGAATTTAAGTAAGAAAGGAACTAACAGTGTGATATT<br>AAGGTGCAATGTTAGACATATAAAACAGTCTTTCACCTCTCTTTGGTTATGTCTTGAATTGGTTTGTTTCTTCACTTATCTGTGTAATCAAGTTTACTATGAGTCTATGATCAAGTAAT<br>TATGCAATCAAGTTAAGTACAGTATAGGCTTT (ZmUbi promoter, OsADH 5’UTR, Cas11 with bpNLS, pea3A terminator) |
| crRNA for *OsPDS* | GGATCATGAACCAACGGCCTGGCTGTATTTGGTGGTTGTGTAGGGAGATGGGGAGAAGAAAAGCCCGATTCTCTTCGCTGTGATGGGCTGGATGCATGCGGGGGAGCGGGAG<br>GCCCAAGTACGTGCACGGTGAGCGGCCCACAGGGCGAGTGAGCGCGAGAGGCGGGAGGAACAGTTTAGTACCACATTGCCCAGCTAACTCGAACGCGACCAACTTATAAA<br>CCCGCGCGCTGTCGCTTGTGTTGTGGATGTGTTGTTTGTGTGATACTATAAAGTTGGTAGATTGTGACTGGCTTAAAAAATCATTAATTAATAATAGGTTATGTTTAGAGTGTTCCC<br>CGCGCCAGCGGGGATAAACCGGTTATGCATGCCTATTGTTGACCATGTTCGCTGTGTTCCCCGCGCCAGCGGGGATAAACCGTTTTTTTGTCCCTTCGAAGGGCAATT (green; OsU6-2 promoter, orange; leader sequences, purple; repeat sequences, red; target sequences for OsPDS, blue; PolyT) |

| | |
|---|---|
| Cas758 | GCCAACATGGTGGAGCACGACACTCTCGTCTACTCCAAGAATATCAAAGATACAGTCTCAGAAGACCAAAGGGCTATTGAGACTTTTCAACAAAGGGTAATATCGGGAAACCTCC<br>TCGGATTCCATTGCCCAGCTATCTGTCACTTCATCAAAAGGACAGTAGAAAAGGAAGGTGGCACCTACAAATGCCATCATTGCGATAAAGGAAAGGCTATCGTTCAAGATGCCTC<br>TGCCGACAGTGGTCCCAAAGATGGACCCCCACCCACGAGGAGCATCGTGGAAAAAGAAGACGTTCCAACCACGTCTTCAAAGCAAGTGGATTGATGTGATAACATGGTGGAGC<br>ACGACACTCTCGTCTACTCCAAGAATATCAAAGATACAGTCTCAGAAGACCAAAGGGCTATTGAGACTTTTCAACAAAGGGTAATATCGGGAAACCTCCTCGGATTCCATTGCCC<br>AGCTATCTGTCACTTCATCAAAAGGACAGTAGAAAAGGAAGGTGGCACCTACAAATGCCATCATTGCGATAAAGGAAAGGCTATCGTTCAAGATGCCTCTGCCGACAGTGGTCC<br>CAAAGATGGACCCCCACCCACGAGGAGCATCGTGGAAAAAGAAGACGTTCCAACCACGTCTTCAAAGCAAGTGGATTGATGTGATATCTCCACTGACGTAAGGGATGACGCACA<br>ATCCCACTATCCTTCGCAAGACCCTTCCTCTATATAAGGAAGTTCATTTCATTTGGAGAGGCCGGTCTAGAGTATTTTTACAACAATTACCAACAACAACAAACAACAAACAACATT<br>ACAATTACTATTTACAATTACAAGCACCATGGGgtaaattctagtttttctccttcattttcttggttaggacccttttctctttttatttttttgagctttgatctttctttaaactgatctattttttaattgattggttatggtgtaaatattacatagctttaactgataa<br>tctgattactttatttcgtgtgtctatgatgatgatgatagttacagAGCCAAGAGGACCGCCGACGGCTCTGAGTTTGAGTCCCCGAAGAAGAAGCGCAAGGTCGAGTCCAACTTCATCAACATCCACGT<br>GCTCATCTCCCACTCGCCGTCCTGCCTCAACAGGGACGACATGAACATGCAGAAGGACGCCATCTTCGGCGGCAAGAGGCGCGTTAGAATTTCCAGCCAGAGCCTCAAGCGC<br>GCCATGAGGAAGTCTGGCTACTACGCCCAGAACATCGGCGAGTCTAGCCTCCGCACAATCCATCTCGCCCAGCTCAGAGATGTGCTCAGGCAAAAGCTCGGCGAGCGCTTCGA<br>CCAGAAGATCATCGATAAGACCCTCGCGCTCCTCAGCGGCAAGTCTGTTGATGAGGCCGAGAAGATCTCCGCCGATGCCGTGACACCATGGGTGGTGGGCGAAATTGCCTGGT<br>TCTGCGAGCAAGTTGCCAAGGCCGAGGCCGACAACCTCGACGACAAGAAGCTCCTCAAGGTCCTCAAAGAGGATATCGCCGCCATCCGCGTGAACCTCCAGCAAGGTGTTGAT<br>ATCGCTCTCTCCGGCAGGATGGCCACCTCTGGCATGATGACAGAGCTTGGCAAAGTGGACGGCGCCATGTCTATCGCCCATGCCATCACCACACACCAGGTGGACTCCGACAT<br>CGACTGGTTCACCGCCGTGGACGATCTCCAAGAACAAGGCTCTGCCCACCTCGGGACCCAAGAATTTTCTTCCGGCGTGTTCTACCGCTACGCCAACATCAACCTCGCGCAGC<br>TCCAAGAGAATCTCGGCGGAGCTTCTAGAGAGCAGGCCCTCGAGATTGCCACACACGTGGTGCATATGCTCGCCACAGAAGTGCCAGGCGCCAAGCAAAGAACCTACGCCGC<br>CTTCAATCCGGCCGACATGGTGATGGTGAACTTCTCCGATATGCCGCTCTCCATGGCCAACGCCTTCGAGAAGGCCGTGAAGGCCAAGGATGGCTTCCTCCAGCCATCCATCC<br>AGGCCTTCAACCAGTATTGGGACCGCGTGGCCAATGGCTACGGCCTTAATGGTGCCGCCGCTCAGTTCTCCCTCTCCGATGTGGATCCAATCACGGCCCAGGTGAAGCAGATG<br>CCGACACTCGAGCAACTCAAGTCCTGGGTGCGCAACAATGGCGAGGCTAAGAGAACAGCCGATGGCAGCGAGTTCGAGAGCCCAAAGAAGAAAAGAAAGGTTGAGGGCTCCG<br>GCGAGGGCAGGGGAAGTCTTCTAACATGCGGGGACGTGGAGGAAAATCCCGGCCCAATGGCCAAGAGGACCGCCGACGGCTCTGAGTTTGAGTCCCCGAAGAAGAAGCGCA<br>AGGTCGAGCGCTCCTACCTCATCCTTAGACTCGCTGGCCCAATGCAGGCTTGGGGCCAGCCAACATTTGAAGGCACCAGACCAACCGGCAGGTTCCCAACAAGATCTGGCCTC<br>CTTGGCCTTCTCGGCGCTTGCCTTGGCATCCAGAGGGATGATACCTCCAGCCTCCAAGCGCTCTCTGAGTCTGTGCAATTCGCCGTGAGGTGCGACGAGCTGATCCTCGATGA<br>TAGACGCGTGTCCGTGACCGGCCTCAGGGATTACCATACAGTTCTTGGCGCTCGCGAGGACTACAGGGGCCTTAAGTCCCACGAGACAATCCAGACCTGGCGCGAGTACCTCT<br>GCGACGCCTCTTTCACAGTTGCCCTCTGGCTCACCCCACACGCCACAATGGTGATCTCCGAGCTTGAGAAGGCCGTGCTCAAGCCGAGGTACACACCTTATCTTGGCAGGCGC<br>AGCTGCCCGCTCACACATCCACTGTTTCTCGGCACATGCCAGGCCTCCGATCCTCAAAAGGCCCTCCTCAACTACGAGCCGGTTGGCGGCGATATCTACTCCGAGGAATCCGT<br>GACAGGCCACCACCTCAAGTTCACCGCTAGGGACGAGCCGATGATCACACTCCCAAGACAATTCGCCAGCCGCGAGTGGTACGTGATCAAAGGCGGCATGGACGTGTCCCAG<br>AAGAGAACAGCTGACGGCAGCGAGTTCGAGAGCCCAAAGAAGAAACGGAAGGTTGAGGGCTCCGGCGAGGGCAGGGGAAGTCTTCTAACATGCGGGGACGTGGAGGAAAAT<br>CCCGGCCCAATGGCCAAGAGGACCGCCGACGGCTCTGAGTTTGAGTCCCCGAAGAAGAAGCGCAAGGTCGAGAACCTCCTCATCGACAACTGGATCCCAGTGAGGCCTAGGA<br>ACGGCGGCAAGGTGCAGATCATCAACCTCCAGAGCCTCTACTGCTCCAGGGATCAATGGCGCCTCTCACTCCCGAGGGACGATATGGAACTCGCTGCTCTCGCTCTCCTCGTG<br>TGCATCGGCCAGATTATTGCCCCAGCCAAGGACGACGTCGAGTTCCGCCATAGGATCATGAACCCGCTCACCGAGGACGAGTTCCAGCAGCTCATTGCCCCGTGGATCGACAT<br>GTTCTACCTCAACCACGCGGAGCACCCGTTCATGCAGACAAAGGGCGTGAAGGCCAACGACGTGACCCCGATGGAAAAGCTCCTTGCTGGCGTTTCCGGCGCCACCAATTGC<br>GCCTTCGTTAACCAGCCAGGCCAAGGCGAAGCTCTTTGCGGAGGCTGTACTGCGATCGCCCTCTTCAACCAGGCCAATCAAGCTCCAGGCTTCGGCGGAGGCTTTAAGTCTGG<br>CCTTAGAGGCGGCACCCCAGTGACCACATTCGTGAGGGGCATTGACCTCAGGTCCACCGTGCTCCTCAACGTGCTCACACTCCCAAGGCTCCAGAAGCAGTTCCCGAACGAGT<br>CCCACACCGAGAACCAGCCGACCTGGATCAAGCCGATCAAGTCCAACGAGAGCATCCCGGCCTCTTCCATCGGCTTCGTTAGGGGCCTCTTCTGGCAGCCAGCGCATATCGA<br>GCTGTGCGATCCAATCGGAATCGGCAAGTGCTCCTGCTGCGGCCAAGAGTCCAACCTCAGGTATACCGGCTTCCTGAAAGAGAAGTTCACCTTCACCGTGAACGGCCTCTGGC<br>CGCATCCACATTCTCCATGCCTCGTGACCGTGAAGAAGGGCGAAGTCGAGGAAAAGTTCCTCGCCTTCACCACAAGCGCCCCAAGCTGGACACAAATCTCCCGCGTGGTGGTG<br>GACAAGATCATCCAGAACGAGAACGGCAACAGGGTCGCCGCCGTGGTGAATCAGTTCCGCAATATCGCCCCACAGAGCCCGCTCGAGCTTATCATGGGCGGCTACCGCAACA<br>ACCAGGCGAGCATTCTTGAGCGCAGGCACGACGTGCTCATGTTCAATCAAGGCTGGCAGCAGTACGGCAACGTGATCAACGAGATCGTCACCGTCGGCCTCGGCTACAAGACA<br>GCCCTTAGAAAGGCCCTCTACACCTTCGCCGAGGGCTTCAAGAACAAGGACTTCAAAGGCGCTGGCGTGAGCGTGCACGAGACAGCTGAGAGGCATTTCTACCGCCAGTCCGA<br>GCTGCTCATCCCAGATGTGCTCGCCAACGTGAACTTCAGCCAGGCGGATGAGGTGATCGCCGACCTCAGGGATAAGCTCCATCAGCTCTGCGAGATGCTGTTCAACCAGAGCG<br>TGGCCCCATACGCGCACCATCCAAAGCTCATCTCTACCCTCGCTCTCGCCAGGGCGACACTCTACAAGCACCTGAGGGAACTCAAGCCGCAAGGCGGCCCATCTAACGGCAA<br>GAGAACTGCTGATGGCTCCGAGTTCGAGAGCCCAAAGAAGAAACGGAAGGTTGAGTGAGAGCTCTGGGCTAATCTAAAACGATTTATCTGTGGCTTCAAGTGTATCGATCACTTA<br>TGTGAGGTGTAATTACTGGTGTTTTTGGTGTGCTCTGGTTCCTTTCAAGTGTGTTGTTGCCGCTCGAACTACTCCGCTATGTAAAACGGTAAAACCTGTTGTCTCATTATGAAAGT<br>GAACTATATTATGTTCTACTACTACTCTACTTAGTCAATTTTCTTCACCTTGATTAGTGTAAATATGAATCTTATATTCTTATGTCTTAAGAAATTAGCACATGTGAAGCCTCCAAGTG<br>CATATTTTCTCGATCGCGAGACGCACAATGCGTGAGAAATTCAGCTGGTTATACTCAAATATATTAATATATCTAGCAGCAGCTCATGGAGATTCAGGAAACTTGGCATCCCTAAT<br>CCCTACCATTTCCATTCTTCCGAGATTGACAGTTCAATACAAGTACAGTAATCTCCTGGTAAGTTTCTTATTAACTTGACATGTAGTAGTAATAATTTGTACGTAGCATAGATACATA<br>GACACAAAAATGTCCTCCCCATTGAGCTAGCCGATTGGAGCCGAACACCGCAGGAATGAATTTACATAATCTGCAAAGAATAAATGGAATGTGCCTCCACAGGAAAACCAGCGG<br>CAGTGTGGCGTTTTCAAGAGCAGCCGTAAGTCGAAGCCTATTCTGAATCGTAGAAATCACTGGGGCATGGTGTAATTACATCCGACTCCAACATCTGTACCACCCTGTGCATTGT<br>AGGCCGCTCCTCTGGCAAAGAACTTACACATTGTTTAGCAAGAGAAAGTAGAGCATCCAAGGTCTCAATCTGCACTCCCTCACAATATGGATCGACAATTTCCCTCTCCCGATTC<br>TCACCGACCAGGAAATTCAACTGCCACAGAGCAAGTAGATTATTTCAAGAATACATTAAATCAATTGAAGGCATACGTAATTCATATCAGAAAACTGTGGATATGAAATGGAAGGA<br>CATAAAGGTCATACATACCCATCCAACAATGTTCAATCCCTTTTCAATAAATGATGCATCAGTAGGTCGTTTTCCGCTTAGTATTTCAAGTAGCAAAACTCCAAAACTGTAGACGTC<br>AGTCTTTTCGGTGGCTCTGCCACTTTGCATATACTC (2x35S promoter, omega, Cas7 with bpNLS (lower and unlderlined letters; catalase intron), T2A, Cas5 with bpNLS, T2A,<br>Cas7 with bpNLS, hsp17.3 terminator) |

| | |
|---|---|
| Cas1163 | TAGAGATAATGAGCATTGCATGTCTAAGTTATAAAAAATTACCACATATTTTTTTGTCACACTTGTTTGAAGTGCAGTTTATCTATCTTTATACATATATTTAAACTTTACTCTACGA<br>ATAATATAATCTATAGTACTACAATAATATCAGTGTTTTAGAGAATCATATAAATGAACAGTTAGACATGGTCTAAAGGACAATTGAGTATTTTGACAACAGGACTCTACAGTTTTA<br>TCTTTTTAGTGTGCATGTGTTCTCCTTTTTTTTTGCAAATAGCTTCACCTATATAATACTTCATCCATTTTATTAGTACATCCATTTAGGGTTTAGGGTTAATGGTTTTTATAGACTAA<br>TTTTTTTAGTACATCTATTTTATTCTATTTTAGCCTCTAAATTAAGAAAACTAAAACTCTATTTTAGTTTTTTTATTTAATAATTTAGATATAAAATAGAATAAAATAAAGTGACTAAAAA<br>TTAAACAAATACCCTTTAAGAAATTAAAAAAACTAAGGAAACATTTTTCTTGTTTCGAGTAGATAATGCCAGCCTGTTAAACGCCGTCGACGAGTCTAACGGACACCAACCAGCG<br>AACCAGCAGCGTCGCGTCGGGCCAAGCGAAGCAGACGGCACGGCATCTCTGTCGCTGCCTCTGGACCCCTCTCGAGAGTTCCGCTCCACCGTTGGACTTGCTCCGCTGTCG<br>GCATCCAGAAATTGCGTGGCGGAGCGGCAGACGTGAGCCGGCACGGCAGGCGGCCTCCTCCTCCTCTCACGGCACGGCAGCTACGGGGGATTCCTTTCCCACCGCTCCTT<br>CGCTTTCCCTTCCTCGCCCGCCGTAATAAATAGACACCCCCTCCACACCCTCTTTCCCCAACCTCGTGTTGTTCGGAGCGCACACACACACAACCAGATCTCCCCCAAATCCA<br>CCCGTCGGCACCTCCGCTTCAAGGTACGCCGCTCGTCCTCCCCCCCCCCCCCTCTCTACCTTCTCTAGATCGGCGTTCCGGTCCATGGTTAGGGCCCGGTAGTTCTACTTCT<br>GTTCATGTTTGTGTTAGATCCGTGTTTGTGTTAGATCCGTGCTGCTAGCGTTCGTACACGGATGCGACCTGTACGTCAGACACGTTCTGATTGCTAACTTGCCAGTGTTTCTCTT<br>TGGGGAATCCTGGGATGGCTCTAGCCGTTCCGCAGACGGGATCGATTTCATGATTTTTTTTGTTTCGTTGCATAGGGTTTGGTTTGCCCTTTTCCTTTATTTCAATATATGCCGT<br>GCACTTGTTTGTCGGGTCATCTTTTCATGCTTTTTTTTGTCTTGGTTGTGATGATGTGGTCTGGTTGGGCGGTCGTTCTAGATCGGAGTAGAATTCTGTTTCAAACTACCTGGTG<br>GATTTATTAATTTTGGATCTGTATGTGTGTGCCATACATATTCATAGTTACGAATTGAAGATGATGGATGGAAATATCGATCTAGGATAGGTATACATGTTGATGCGGGTTTTACT<br>GATGCATATACAGAGATGCTTTTTGTTCGCTTGGTTGTGATGATGTGGTGTGGTTGGGCGGTCGTTCATTCGTTCTAGATCGGAGTAGAATACTGTTTCAAACTACCTGGTGTAT<br>TTATTAATTTTGGAACTGTATGTGTGTGTCATACATCTTCATAGTTACGAGTTTAAGATGGATGGAAATATCGATCTAGGATAGGTATACATGTTGATGTGGGTTTTACTGATGCAT<br>ATACATGATGGCATATGCAGCATCTATTCATATGCTCTAACCTTGAGTACCTATCTATTATAATAAACAAGTATGTTTTATAATTATTTTGATCTTGATATACTTGGATGATGGCATA<br>TGCAGCAGCTATATGTGGATTTTTTTAGCCCTGCCTTCATACGCTATTTATTTGCTTGGTACTGTTTCTTTTGTCGATGCTCACCCTGTTGTTTGGTGTTACTTCTGCAGGTCGAC<br>TCTAGAGAATTCCAAGCAACGAACTGCGAGTGATTCAAGAAAAAAGAAAACCTGAGCTTTCGATCTCTACGGAGTGGTTTCTTGTTCTTTGAAAAAGAGGGGGATTAACTAGAT<br>GGCCAAGAGGACCGCCGACGGCTCTGAGTTTGAGTCCCCGAAGAAGAAGCGCAAGGTCGAGGCCGACGAGATCGATGCCATGGCTCTTTACAGAGCGTGGCAGCAGCTCG<br>ACAATGGCTCTTGCGCTCAGATTCGCAGGGTGTCCGAGCCAGATGAGCTGAGGGATATTCCGGCCTTCTACAGGCTCGTGCAGCCATTCGGCTGGGAGAACCCAAGACATCA<br>GCAGGCGCTCCTCCGCATGGTTTTCTGCCTTTCCGCCGGCAAGAACGTGATCCGCCACCAGGATAAGAAGTCCGAGCAGACCACCGGCATTTCTCTCGGCAGAGCCCTTGCT<br>AACTCCGGCAGGATCAATGAGCGCCGCATCTTCCAGCTCATCAGGGCCGATAGGACAGCCGACATGGTTCAGCTCAGACGCCTCCTCACACATGCCGAGCCAGTTCTCGATT<br>GGCCACTCATGGCCAGGATGCTCACATGGTGGGGAAAGAGAGAGCGCCAGCAGCTCCTTGAGGACTTCGTGCTCACCACCAACAAGAACGCCAAGCGCACAGCTGACGGCA<br>GCGAGTTTGAGAGCCCAAAGAAAAAGAGGAAGGTCGAAGGCTCCGGCGAGGGCAGGGGAAGTCTTCTAACATGCGGGGACGTGGAGGAAAATCCCGGCCCAATGGCCAAG<br>AGGACCGCCGACGGCTCTGAGTTTGAGTCCCCGAAGAAGAAGCGCAAGGTCGAGTACCTCTCCAAGGTGATCATTGCCCGCGCTTGGTCCAGGGACCTCTACCAACTTCATC<br>AAGGCCTCTGGCATCTCTTCCCGAATCGCCCTGATGCCGCCAGGGATTTCCTGTTCCATGTCGAGAAGAGGAACACCCCAGAGGGCTGCCATGTGCTCCTCCAATCTGCCCA<br>AATGCCAGTGTCTACCGCCGTGGCCACCGTGATCAAGACCAAGCAGGTTGAGTTCCAGCTCCAGGTGGGCGTGCCACTCTATTTCAGGCTCAGGGCGAACCCGATCAAGACG<br>ATCCTCGACAACCAGAAGCGCCTCGACTCCAAGGGCAACATCAAGAGGTGCAGGGTGCCGCTCATCAAAGAGGCCGAGCAAATTGCCTGGCTCCAGCGCAAGCTTGGCAAT<br>GCTGCCAGAGTCGAGGATGTGCACCCAATCTCTGAGCGCCCGCAATACTTCTCTGGCGACGGCAAGTCTGGCAAGATCCAGACCGTTTGCTTCGAGGGCGTGCTCACCATCA<br>ATGATGCCCCAGCGCTCATCGACCTCGTGCAGCAAGGCATTGGACCGGCTAAGTCTATGGGCTGCGGCCTCCTCTCACTCGCTCCACTTAAGAGAACAGCGGACGGCAGCG<br>AGTTCGAGAGCCCAAAGAAGAAAAGAAAGGTTGAGGGCTCCGGCGAGGGCAGGGGAAGTCTTCTAACATGCGGGGACGTGGAGGAAAATCCCGGCCCAATGGCCAAGAGG<br>ACCGCCGACGGCTCTGAGTTTGAGTCCCCGAAGAAGAAGCGCAAGGTCGAGGAACCGTTCAAGTACATCTGCCACTACTGGGGCAAGTCCTCCAAGAGCCTCACCAAGGGC<br>AACGATATCCACCTCCTCATCTACCACTGCCTCGATGTTGCCGCCGTTGCCGATTGCTGGTGGGATCAATCTGTGGTGCTCCAGAACACCTTCTGCCGCAACGAGATGCTCTC<br>CAAGCAACGCGTTAAGGCCTGGCTCCTCTTCTTCATTGCCCTCCACGACATCGGCAAGTTCGACATCCGCTTCCAGTACAAGTCCGCCGAGTCCTGGCTCAAGCTCAACCCA<br>GCTACACCAAGCCTCAACGGCCCGTCCACACAAATGTGCCGCAAGTTCAATCATGGCGCCGCTGGCCTCTACTGGTTCAACCAGGATTCTCTCAGCGAGCAGTCCCTCGGCG<br>ACTTCTTCTCCTTCTTCGATGCCGCGCCACATCCGTACGAGAGCTGGTTCCCATGGGTTGAAGCCGTGACAGGCCACCATGGCTTCATCCTCCATAGCCAGGACCAGGACAA<br>GTCCAGGTGGGAGATGCCAGCTTCTCTCGCCTCTTACGCCGCGCAGGATAAGCAAGCTCGCGAGGAATGGATCTCCGTGCTCGAGGCTCTGTTCCTCACACCAGCCGGCCT<br>CTCCATCAACGACATTCCGCCAGATTGCTCCTCTCTCCTCGCTGGCTTTTGCTCCCTCGCTGATTGGCTTGGCTCCTGGACCACCACCAACACCTTTCTGTTCAACGAGGACG<br>CCCCGTCCGACATCAATGCCCTCAGGACCTACTTCCAGGACAGGCAGCAAGATGCCTCCAGGGTGCTCGAACTCTCTGGCCTCGTTTCCAACAAGCGCTGCTACGAGGGTGT<br>GCATGCGCTCCTCGATAACGGCTACCAGCCAAGACAGCTCCAGGTCCTCGTGGATGCCCTTCCAGTTGCTCCAGGCCTCACAGTGATTGAGGCTCCAACAGGCTCCGGCAAG<br>ACAGAGACAGCCCTCGCCTATGCCTGGAAGCTCATCGATCAGCAGATCGCCGACTCCGTGATCTTCGCGCTCCCAACACAAGCCACCGCCAATGCCATGCTCACCAGGATGG<br>AAGCCTCTGCCAGCCACCTCTTCAGCTCCCCGAATCTCATTCTCGCCCACGGCAACTCCCGCTTCAACCACCTCTTTCAGTCCATCAAGTCCCGCGCCATCACCGAGCAAGG<br>CCAAGAAGAGGCTTGGGTGCAGTGCTGCCAATGGCTCTCCCAGTCCAACAAGAAGGTTTTCCTCGGCCAGATCGGCGTGTGCACCATTGACCAGGTGCTCATCTCTGTGCTC<br>CCGGTGAAGCACAGATTCATTCGCGGCCTTGGCATCGGCCGCTCAGTGCTCATTGTGGATGAGGTGCACGCCTACGATACCTACATGAACGGCCTTCTCGAGGCCGTGCTCA<br>AGGCCCAAGCCGATGTTGGCGGCTCTGTGATCCTCCTCTCTGCGACACTCCCAATGAAGCAGAAGCAAAAGCTCCTCGACACCTACGGCCTCCACACCGATCCAGTCGAGAA<br>CAACTCCGCCTATCCGCTCATCAATTGGAGGGGCGTTAACGGCGCCCAGAGGTTCGATCTCCTTGCGCATCCAGAGCAGCTCCCGCCAAGGTTCTCTATCCAGCCTGAGCCA<br>ATCTGCCTCGCCGACATGCTCCCAGATCTCACCATGCTCGAGCGCATGATCGCCGCTGCTAATGCCGGTGCTCAAGTGTGCCTCATCTGCAACCTCGTCGATGTGGCCCAGG<br>TGTGCTACCAGAGGCTCAAAGAGCTGAACAACACCCAGGTGGACATCGACCTCTTCCACGCCAGGTTCACCCTCAACGACCGCCGCGAGAAAGAGAACCGCGTCATCTCCAA<br>CTTCGGCAAGAACGGCAAGAGGAACGTGGGCAGAATCCTCGTGGCCACACAGGTGGTGGAACAGTCTCTCGACGTGGACTTCGACTGGCTCATCACCCAACACTGCCCAGC<br>CGACCTGCTCTTTCAAAGACTCGGCAGGCTCCACCGCCACCACAGAAAGTACAGACCAGCGGGCTTCGAGATCCCGGTCGCCACAATTTTGCTGCCGGATGGCGAAGGCTA<br>CGGCAGGCACGAGCATATCTACTCCAACGTGAGGGTCATGTGGCGCACCCAGCAGCACATCGAGGAACTCAATGGCGCCAGCCTGTTCTTCCCGGATGCCTACAGACAGTG<br>GCTCGACTCCATCTACGACGACGCCGAGATGGATGAGCCTGAGTGGGTTGGCAACGGCATGGACAAGTTCGAGTCTGCCGAGTGCGAGAAGAGGTTCAAGGCTCGCAAGGT<br>GCTGCAGTGGGCCGAAGAGTACTCCCTCCAGGACAACGACGAGACAATCCTCGCCGTGACACGCGACGGCGAAATGTCTCTCCCACTCCTTCCGTACGTGCAGACCTCTTCT<br>GGCAAGCAACTCCTCGACGGCCAGGTGTACGAGGATCTCTCTCATGAGCAGCAGTACGAGGCCCTCGCGCTCAATAGAGTGAACGTGCCGTTTACCTGGAAGCGCTCCTTCA<br>GCGAGGTGGTCGATGAAGATGGCCTCCTCTGGCTCGAGGGCAAGCAGAATCTTGATGGCTGGGTGTGGCAGGGCAACTCCATCGTGATCACCTACACAGGCGACGAGGGCA<br>TGACCAGGGTGATCCCAGCCAATCCAAAGAAGAGAACGGCGGACGGCAGCGAGTTCGAGAGCCCAAAGAAAAAGCGGAAGGTTGAGTGAGAGCTCCGATCCAGGCCTCCCA<br>GCTTTCGTCCGTATCATCGGTTTCGACAACGTTCGTCAAGTTCAATGCATCAGTTTCATTGCCCACACACCAGAATCCTACTAAGTTTGAGTATTATGGCATTGGAAAAGCTGTT<br>TTCTTCTATCATTTGTTCTGCTTGTAATTTACTGTGTTCTTTCAGTTTTTGTTTTCGGACATCAAAATGCAAATGGATGGATAAGAGTTAATAAATGATATGGTCCTTTTGTTCATTC<br>TCAAATTATTATTATCTGTTGTTTTTACTTTAATGGGTTGAATTTAAGTAAGAAAGGAACTAACAGTGTGATATTAAGGTGCAATGTTAGACATATAAAACAGTCTTTCACCTCTCTT<br>TGGTTATGTCTTGAATTGGTTTGTTTCTTCACTTATCTGTGTAATCAAGTTTACTATGAGTCTATGATCAAGTAATTATGCAATCAAGTTAAGTACAGTATAGGCTTT(ZmUbi<br>promoter, OsADH 5'UTR, Cas11 with bpNLS, T2A, Cas6 with bpNLS, T2A, Cas3 with bpNLS, pea3A terminator) |
